# Attention-dependent attribute comparisons underlie multi-attribute decision-making in orbitofrontal cortex

**DOI:** 10.1101/2024.11.12.623291

**Authors:** Aster Q. Perkins, Erin L. Rich

## Abstract

Economic decisions often require weighing multiple dimensions, or attributes. The orbitofrontal cortex FC) is thought to be important for computing the integrated value of an option from its attributes and comparing values to make a choice. Although OFC neurons are known to encode integrated values, evidence for value comparison has been limited. Here, we used a multi-attribute choice task for monkeys to investigate how OFC neurons integrate and compare multi-attribute options. Attributes were represented separately and eye tracking was used to measure attention. We found that OFC neurons encode the value of attended attributes, independent of other attributes in the same option. Encoding was negatively weighted by the value of the same tribute in the other option, consistent with a comparison between the two like attributes. These results indicate that OFC computes comparisons among attributes rather than integrated values, and does so dynamically, shifting with the focus of attention.

## Introduction

We are routinely faced with choices that involve multiple dimensions, such as cost, risk, quality, or quantity. To make a decision, it is thought that the brain first combines relevant features to compute the overall value, or utility, of each option, then compares these values to arrive at a choice (*1–5*). The orbitofrontal cortex (OFC) is critical for value-based decision-making (*6–9*), and has been proposed to carry out this process of value integration and comparison (*3, 10–12*). Consistent with this, neural activity in OFC correlates with integrated values (*3, 9–11, 13–17*), but so far there has been scant evidence for comparison of these values in OFC ((*14, 18–21*) but see (*9, 22*)). In contrast, comparison signals have been found in other brain regions, including the neighboring ventromedial PFC (vmPFC), ventral striatum, and amygdala (*4, 23–27*). These comparison signals consist of antagonism between values of each option, effectively computing a weighted value difference. The relative ambiguity of comparison signals in OFC has contributed to speculation that it is not the site of choice computation per se, but plays other roles in decision-making such as credit assignment (*28, 29*), learning (*30, 31*), or representing a relational map of task space (*32, 33*).

An often-overlooked feature of OFC is that many neurons encode the value of the individual dimensions, or attributes, of a choice option (*9, 13–15, 18, 34, 35*). These responses are understudied compared to the more prevalent integrated value signals, making it unclear whether comparisons might be computed among unintegrated attributes. Behavioral evidence supports this idea, showing that decision-makers often compare component attributes directly, for instance cost versus cost or taste versus taste, either instead of or in addition to comparing integrated values (19–24). Similarly, eye tracking in humans and monkeys has found that gaze frequently shifts between like attributes of the two options, consistent with a decision process that involves comparison of similar attributes (*36, 37*).

Shifts of visual attention not only provide insights to the decision process, but affect value-related responses in OFC. In the absence of choice, directing attention to a reward-predicting cue amplifies OFC encoding of the cue’s value (*28, 29*). Similarly, when choice options are presented sequentially, many neurons code the value of the currently presented option rather than having selectivity for only one (*9, 18, 19, 22, 38*). With this in mind, we aimed to test whether neurons in OFC encode and compare component attributes of multi-attribute options, and how this is affected by natural shifts of visual attention in the course of making a decision. To do this, we designed a novel multi-attribute choice task for rhesus monkeys, in which attribute values were represented by separate visual cues that were presented simultaneously and varied independently. Using gaze to measure visual attention, we show that single neurons in OFC encode the value of attended attributes independent of other attributes in the same option. Moreover, this encoding is negatively weighted by the value of the same attribute in the other option, consistent with a comparison between like attributes. These results extend previous work showing that visual attention is a critical mediator of neural activity in OFC (*22, 39, 40*), and identify a novel, attention-based mechanism of choice computation at the level of single neuron activity.

## Results

### Choice behavior indicates attribute-level comparisons

Two rhesus monkeys performed a multi-attribute decision-making task (**Fig 1a**). Choice options varied in sweetness (sucrose concentration) of a fluid reward and probability of delivery. The sweetness and probability of an option were each represented by a colored bar, whose height varied monotonically with attribute magnitude. Bar color indicated whether the mapping between height and value was direct (i.e., bigger is better) or indirect (bigger is worse). Sweeter and more probable outcomes were indicated by larger blue and yellow bars (direct mapping) or smaller magenta and green bars (indirect mapping) (**Fig 1b**).

**Figure 1.**
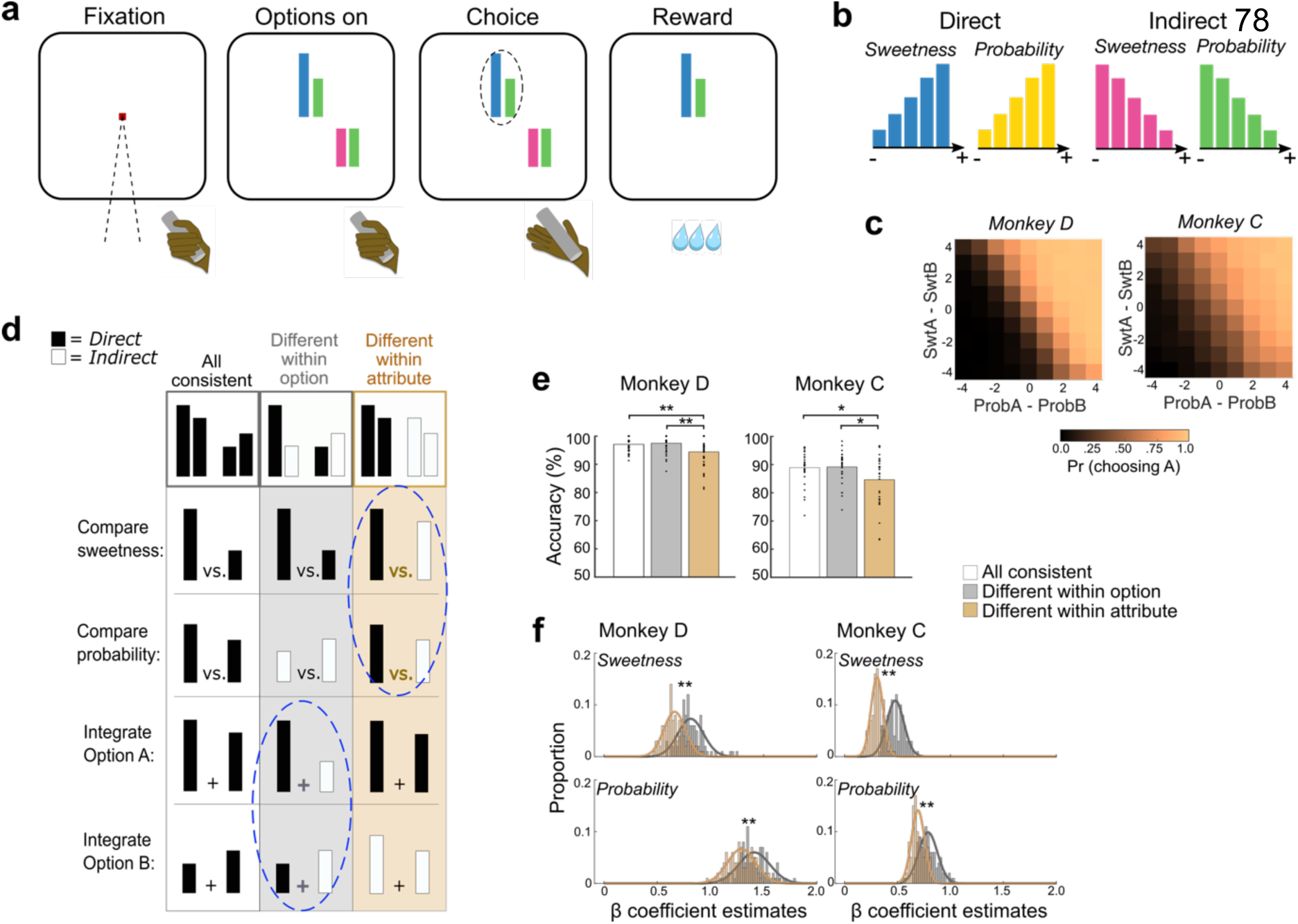
Multi-attribute choice task. (**a**) Left to right: Monkeys initiated a trial by holding a touch-sensitive bar and fixating a central point. Choice options were represented by a pair of colored bars of variable height. The left bar indicated the sweetness of an option and the right bar indicated the probability of receiving it. Monkeys freely viewed the options while holding the touch bar, and reported a choice by releasing the bar while directing gaze to one option. (**b**) Sweetness and probability attributes were represented by colored bars, whose height indicated the value of the attribute. Values varied monotonically in 5 increments. Blue and yellow bar heights directly mapped to the value of the attribute (-= low, + = high). Magenta and green bar heights indirectly mapped to the value of the attribute. (**c**) The probability of choosing option A (arbitrarily designated) increased with the difference between option A and B probabilities (ProbA – ProbB) and sweetnesses (SwtA – SwtB). Differences were computed from ordinal levels of each attribute. (**d**) Schematic of three trial types. Top row: Example decisions between two pairs of bars. White column: all attributes have the same mapping, either all direct (shown) or all indirect (not shown). All putative choice computations would take place between bars of the same mapping. Gray: mappings differ among attributes within an option, but like attributes are the same, so that attribute comparisons take place between bars with the same mapping, but integration takes place across different mappings (blue dashed circle). Orange: mappings differ within like attributes but are the same within an option, so that attribute comparisons take place between bars with different mappings (blue dashed circle), but value integration takes place between bars with the same mapping. (**e**) Accuracy on trials with an objectively better option is consistently lower when like attributes differ (orange). Bars show averages, points show session-wise accuracy in each trial type. Post-hoc comparisons *p≤0.05, **p≤0.01. (**f**) Weights for each attribute were estimated by beta coefficients from logistic regressions predicting choice probabilities separately from groups of trials in which attribute mapping differed within option (gray) or within attribute (orange) as in **d**. Since two coefficients of opposite sign were estimated for each attribute (for option A and B), plots show the mean absolute value of these. Histograms show bootstrapped sets of trials of each type. Lower coefficients indicate more choice variability. ** Wilcoxon rank-sum test p<0.001.

Monkeys D and C performed 43 and 30 sessions respectively, and averaged 1,085 and 1,203 trials per session. Choice probabilities indicated that they weighted both attributes to select sweeter, more probable options (**Fig. 1c**). Foundational models of economic choice, such as expected utility theory (*41*) and prospect theory (*42*) assume that probability is multiplicatively combined with other attributes to calculate option values (but see (*43, 44*)). In contrast, we previously found that behavior in this task is better fit by models that combine attributes linearly (*37*). Formal model comparisons found that choice behavior in this data set was also better fit by an additive rather than multiplicative model (**Fig S1a**). Since these models are highly similar, this was confirmed with model recovery, showing that we can reliably select the correct generative model from simulated choices (**Fig S1b**). Therefore, choice behavior of both monkeys indicated that they computed choices by weighing and linearly combining sweetness and probability attributes, and we used this model for further analyses (**Fig S1c-j**).

Next, we separated trials with different attribute mappings to assess inefficiencies in the decision process (*37*). We considered two operations that could be involved in computing decisions: integrating values within an option, and directly comparing like attributes of the different options (i.e., sweetness vs. sweetness and probability vs. probability). Integration or comparison should be easier when the relevant bars have the same mapping, since combining the area of two bars or deciding which of two bars is taller could be performed with simple perceptual judgements. However, when the relevant bars have different mappings (direct vs. indirect), the monkeys must translate across mappings by referencing an internal representation of the bar’s meaning – either its position on a value scale or its specific attribute magnitude (e.g., 50% probability) – in order to integrate or compare the information. This process can introduce subtle but measurable inefficiencies into the choice process (*37*). To measure this, we separately analyzed behavior during three trial types. First were trials in which all four bars had a consistent mapping (all direct or all indirect, **Fig 1d**, white), so that integration of attributes within an option or comparison of like attributes across options would always rely on bars with the same mapping. We compared these to trials in which two bars were direct and two were indirect, in different arrangements. When mappings differed within option but like attributes had the same mapping (i.e., sweetness matched sweetness and probability matched probability), then attribute-level comparisons would take place between bars with the same mapping, but value integration would involve bars with different mappings (**Fig 1d**, gray). On the other hand, if the two attributes within each option had the same mapping but differed from the two attributes of the other option, then combining sweetness and probability into integrated values would involve bars of the same mapping, whereas comparing like attributes without integration would occur across different mappings (**Fig 1d**, orange). Therefore, separating trials according to whether mappings are mismatched within option or within attribute allowed us to test whether the need to translate between mappings impeded specific component processes that might underlie choice computation.

We first tested this without relying on behavior models by subselecting trials in which one option was superior to the other in both sweetness and probability, and therefore was objectively better (n = 12,038 and 8,888 trials across all sessions, Monkey D and C respectively). Focusing on these trials allowed us to compute choice accuracies needing to account for subjective weighting of sweetness versus probability. In both subjects, accuracy on choices with an objectively better option was lower when mappings differed within like attributes, but was unaffected when mappings differed within an option (ANOVA Monkey D: F_2,126_=9.32, p=1.68×10^−4^, Tukey-corrected post-hoc comparisons: all consistent vs. differ within option p=0.86, all consistent vs differ within attribute p=0.002, differ within option vs differ within attribute p=2.13×10^−4^; Monkey C: F_2,126_=9.32, p=1.68×10^−4^, Tukey-corrected post-hoc comparisons: all consistent vs differ within option p=0.99, all consistent vs differ within attribute p=0.050, differ within option vs differ within attribute p=0.036) (**Fig 1e**). Consistent with our previous findings (*37*), this pattern suggests that choices in this task involve, at least in part, comparisons between like attributes.

Next, we used the same attribute arrangements to test for attribute comparisons across all choices, not just those with an objectively better option. To do this, we separately modeled trials in which bar mappings differed within an option (**Fig 1d**, gray) or within like attributes (**Fig 1d**, orange) with logistic regressions (*Eq 1,3*). If different bar mappings impeded within-attribute comparisons, we would expect less consistent choices on trials where mappings differed within like attributes (orange), quantified as shallower fitted slopes. Indeed, estimated beta coefficients were consistently lower on these trials, indicating more choice variability (**Fig 1f**). Taken together, patterns of choice behavior provide consistent evidence that attribute comparisons contribute to decisions in this task. Given this, our goal was to assess whether and how such comparisons are made by OFC neurons.

### OFC neurons encode attributes of multi-attribute options

We recorded a total of 322 well-isolated single neurons from OFC as monkeys performed the multi-attribute choice task (163 Monkey D, 159 Monkey C). To understand how neurons encoded information about sweetness and probability, we analyzed each neuron’s activity in sliding windows with a regression model predicting firing rate from the value of each attribute of the chosen and unchosen options, as well as the mappings (direct or indirect) of each attribute (*Eq. 5*). Neurons that encode the integrated value of a choice should have non-zero regression coefficients for both attributes of the chosen option, simultaneously with the same sign. Some neurons fit these criteria (**Fig 2a**), but this encoding pattern was relatively infrequent and accounted for less than 10% of neurons at any time (**Fig 2b, S2e**). On the other hand, neurons that compute comparisons between chosen and unchosen attributes should encode both values with opposite signs, but this pattern was even less common (**Fig 2b, S2e**). Instead, many neurons encoded the value of only one attribute, most frequently an attribute of the chosen option (**Fig 2a, S2b**). Single attribute responses were found in approximately 30% of neurons in any time bin during the choice epoch, making them the most common type of response (**Fig 2b, S2e**).

**Figure 2.**
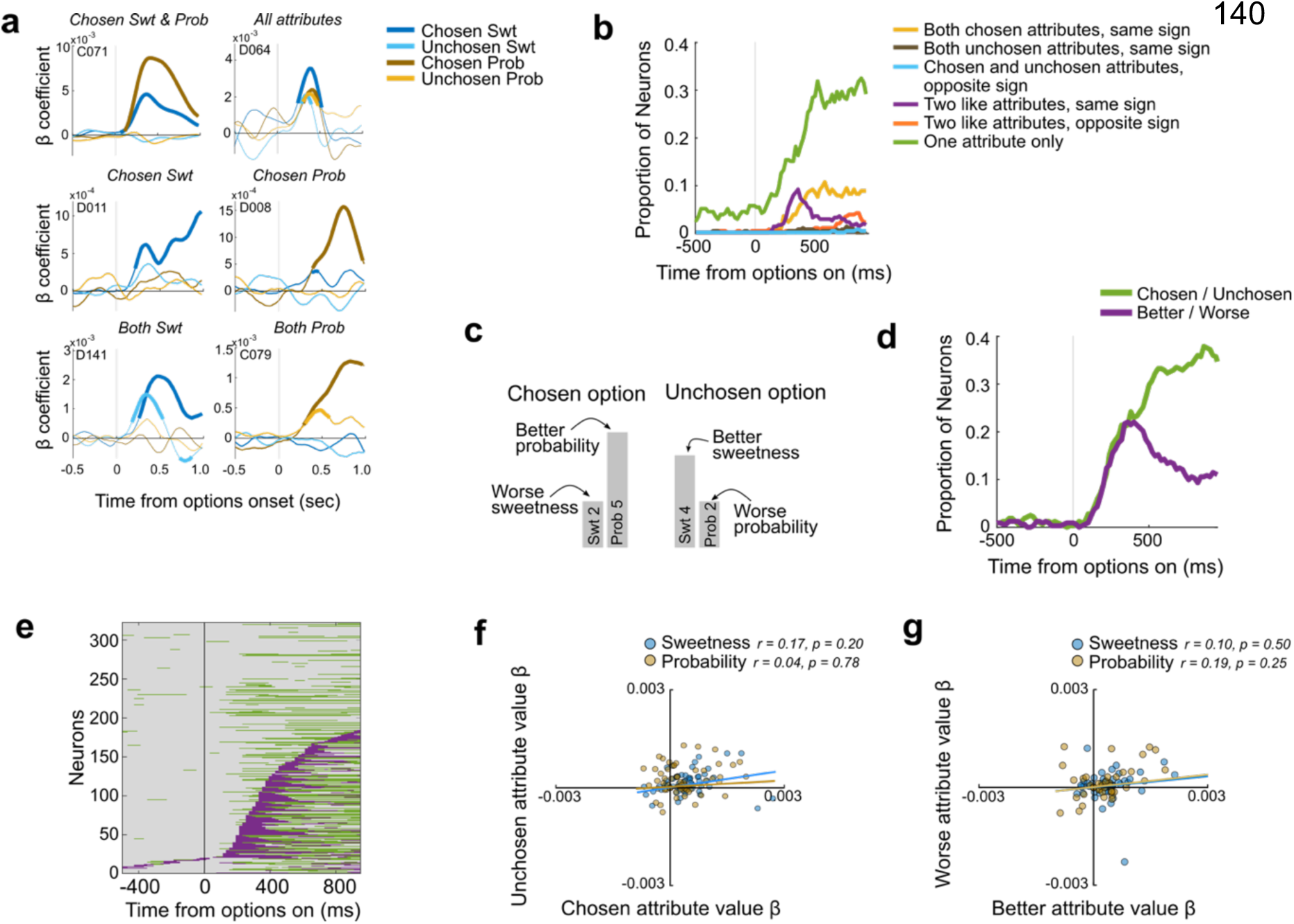
Attributes encoded by OFC neurons. (**a**) Types of attribute encoding in six example neurons. Lines show signed coefficient estimates for chosen or unchosen sweetness (Swt) or probability (Prob) over time. Thick segments denote regions of significance (p≤0.01 x 3 consecutive time bins). Top row left to right: A neuron encoding both sweetness and probability of the chosen option. A neuron whose firing rate varies with all four attributes. Middle row: Neurons encoding the chosen sweetness only (left) or chosen probability only (right). Bottom row: Neurons whose firing rates vary with both chosen and unchosen sweetnesses (left) or probabilities (right). (**b**) Proportion of neurons with different encoding patterns. (**c**) Schematic showing relative value of an attribute that is discordant with the chosen options. The choice is between a low (level 2) sweetness and high (level 5) probability versus a high (level 4) sweetness and low (level 2) probability, and the first option is selected. Only direct mappings are shown for simplicity. The chosen/unchosen model would assign attribute values for chosen sweetness, chosen probability, unchosen sweetness, unchosen probability as [2 5 4 2], but the better/worse model for the same trial would assign better sweetness, better probability, worse sweetness, worse probability as [4 5 2 2]. (**d**) Proportion of neurons assigned to each model. (**e**) Neurons assigned to the best/worst (purple) or chosen/unchosen (green) model over time, sorted by the time of better/worse encoding. Many neurons shifted from encoding better/worse attributes early and chosen/unchosen attributes later. (**f-g**) There were no correlations between beta coefficients for chosen and unchosen attributes or better and worse attributes.

We also found neurons that encoded the same attribute of both chosen and unchosen options, with firing rates modulated in the same direction (e.g., higher firing with higher value of either attribute) (**Fig 2a, S2c**). This type of response is inconsistent with a comparison signal, but was found in ∼10% of neurons, soon after the options appeared (**Fig 2b, S2e**). We reasoned that, if these neurons encode unintegrated attributes before a choice is determined, categorizing attributes according to which option was chosen or not chosen would be the wrong reference frame for quantifying these responses. Instead, the neurons might distinguish attributes according to whether they are *better* or *worse* than the other like attribute on that trial, regardless of which one belongs to the option that is ultimately chosen (**Fig 2c**). Therefore, we tested whether neurons encoded the values of the better or worse sweetness or probability (*Eq. 6*). Since better attributes tend to be chosen, this model is correlated with the previous one, so we used model comparisons to distinguish them.

We found that up to 20% of OFC neurons were best fit by the better/worse model, indicating that they encoded like attributes relative to one another, not relative to the choice (**Fig 2d**). This encoding was most frequent early in the choice, and about 500ms later gave way to a predominant encoding of attributes according to whether they were chosen or not. This pattern suggests that OFC initially processes unintegrated attributes in a reference frame that does not depend on, and therefore can precede, a choice. We considered additional models in which attributes of an option are integrated in different ways, but the most common encoding in both subjects was of independent attributes, either as the better or worse sweetness/probability early in the trial, or as part of the chosen or unchosen option later in the trial (**Fig S2f**). There was a tendency for the same neurons to shift from coding attributes relative to one another (better/worse reference frame) to encoding them relative to the choice (chosen/unchosen reference frame) (binomial test, p = 5.15×10^−4^), although there were also many neurons that encoded chosen/unchosen attributes without previously encoding better/worse attributes (**Fig 2e**). This pattern was found in each monkey separately, but only significant in one (Monkey D, C: p = 0.002, p =0.056).

So far, we found that many OFC neurons encode attributes separately. Early in the choice, they are encoded in relation to the same attribute in the other option (i.e., better/worse reference frame) and later in relation to the decision (i.e., chosen/unchosen reference frame). This could suggest a comparison process between better and worse attributes that produces a chosen option. Therefore, we tested for antagonistic value coding consistent with a comparison signal, by looking for inverse relationships among the regression coefficients. However, in both the better/worse reference frame and chosen/unchosen reference frame, we found no significant relationships (**Fig 2f-g**). Therefore, although encoding patterns are suggestive of a transition from encoding attributes in a pre-choice to post-choice frame of reference, this analysis did not find evidence for within-attribute comparisons in OFC.

### Gaze behavior relates to attributes and attribute values

Since our task represented attributes with separate cues, we were able to use eye tracking to assess visual exploration of attributes and whether directing gaze to an attribute affects neural responses in OFC. To do this, we defined pre-choice fixations as those directed to an attribute bar between the time of option presentation and choice (**Fig 3a**). Since monkeys had to fixate an option to report their choice, we excluded this final fixation that coincided with the bar release. Both monkeys visually explored the choice options, but often did not foveate all four attributes on the screen (**Fig 3b**). Since both attributes contributed to their choices, this suggests that subjects used both directed gaze and peripheral vision to evaluate the options. Consistent with this, the first fixation of both monkeys was more likely to be on the higher value option on objective trials, where both attributes of one option were more valuable than both attributes of the other (**Fig 3c**). Monkey D and C fixated the higher value option on 76.5% and 68.5% of these trials, suggesting that they rapidly evaluated peripheral targets before shifting their gaze, as monkeys do on single attribute choices (*45*). However, our subjects were less proficient at this covert evaluation when attributes needed to be weighed. Across all trials, including those in which the higher value sweetness and higher value probability belong to different options, their tendencies to look first at the chosen option dropped to 62.2% and 52.7% (Monkey D, C). If the subjects evaluated unintegrated attributes, however, we might expect their first fixations to be directed to better attributes, regardless of which option was chosen. Indeed, both subjects’ first fixations were directed to the better sweetness or probability on each trial more consistently than to the chosen option (**Fig 3d**). This tendency was stronger for sweetnesses, but also present for probability (**Fig 3e**). Taken together, patterns of gaze behavior indicate that subjects used peripheral vision to evaluate unintegrated attributes, and they were biased to look at valuable attributes first as they made their choices.

**Figure 3.**
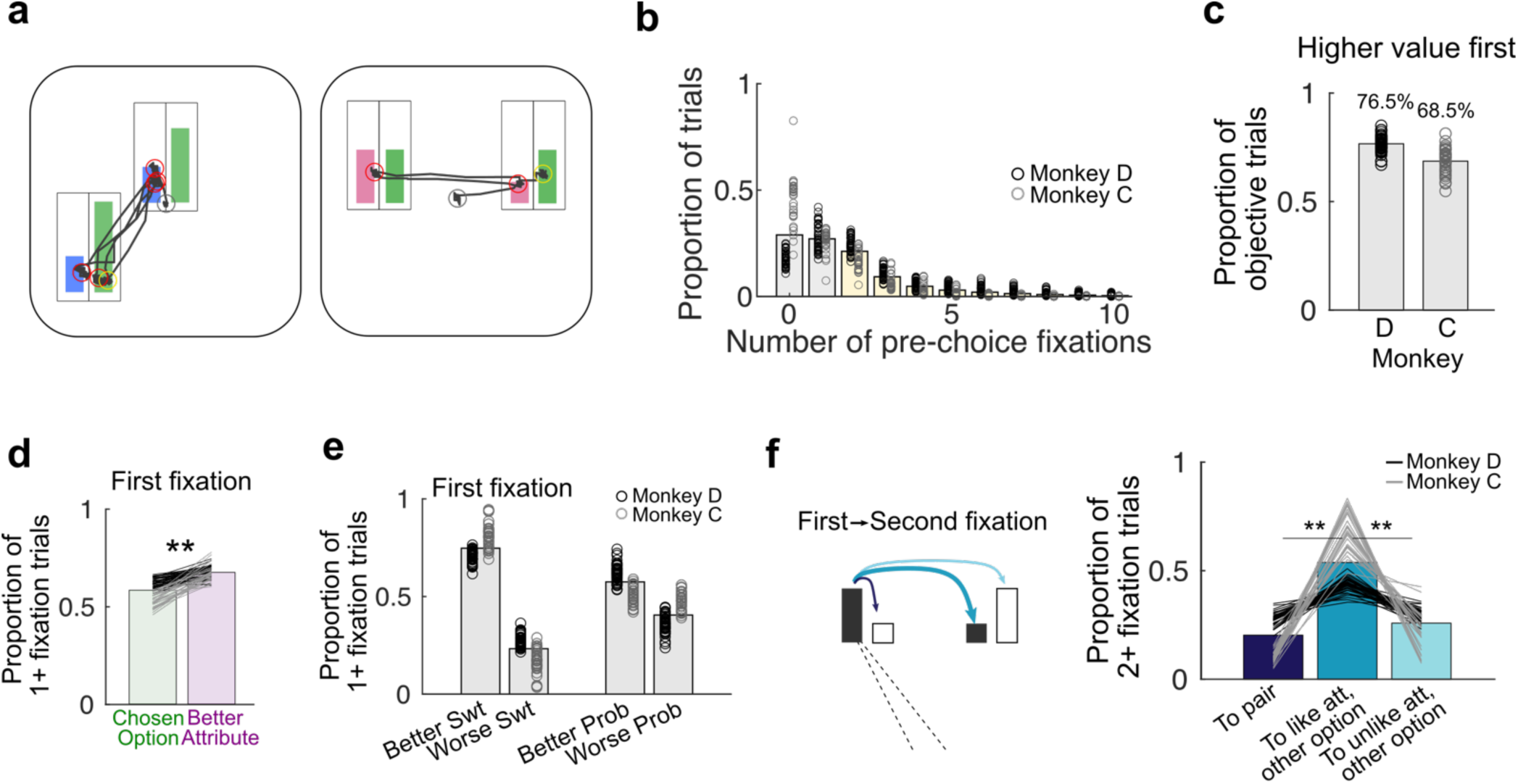
Fixation patterns during multi-attribute choices. (**a**) Gaze paths (black) on two example trials. Outlines around the colored bars show inclusion regions assigned to each attribute (not visible to the monkey). Red circles indicate fixations included in analyses. Fixations on the central fixation point at trial start (gray circle) and on the selected option at bar release (yellow) were excluded. (**b**) Number of pre-choice fixations made by each monkey. Bars = average across monkeys and sessions, circles = session averages. Yellow bars = trials included in the gaze-aligned neural analyses. (**c**) Proportion of trials in which both attributes of one option were higher value than both attributes of the other, and the monkey looked at the higher value option first. Bars = average across sessions, circles = average in each session. (**d**) Proportion of trials with one or more pre-choice fixation in which the monkey first looked at an attribute in the option he would ultimately choose (green) or at an attribute that was higher value than the like attribute of the other option, regardless of choice (purple). Bars = average across monkeys and sessions, lines = session averages. Black = Monkey D: Gray = Monkey C. Proportions were compared with sign-rank test across sessions. ** Monkeys D: p=1.59×10^−8^, C: p=1.73×10^−6^. (**e**) Proportion of trials in which the monkey first looked at a sweetness bar (Swt) and it was the better or worse sweetness (left), or first looked at a probability bar (Prob) and it was the better or worse probability (right). Bars = average across sessions and monkeys, circles = session proportions. (**f**) Schematic of three types of gaze transitions in an example where the first fixation is on the sweetness bar of the left option (left), and proportions of each type of gaze transition (right). Navy: to the paired attribute; Dark teal: to the like attribute of the other option; light teal: to the unlike attribute of the other option. Bar plots = average across sessions and monkeys, lines = proportions within sessions. One-way ANOVA of session-wise measures: Monkeys D: F_2,128_=71.4, p=1.86×10^−21^, C: F_2,89_=65.5, p=4.50×10^−18^. ** Bonferroni-corrected post-hoc comparisons p<0.001.

If the monkeys made choices by comparing like attributes, they might also sequentially evaluate the same attribute of each option. However, if their decision strategy relied on integrating the values of two attributes within an option, they might sequentially evaluate the sweetness and probability bar of a single option. To assess these possibilities, we categorized the first gaze transition on trials with at least 2 pre-choice fixations as shifting either to the paired attribute within the same option (e.g., sweetness A to probability A), to the like attribute of the other option (e.g., sweetness A to sweetness B), or to the nonmatching attribute in the other option (e.g., sweetness A to probability B) (**Fig 3f**). We found that gaze more frequently shifted between like attributes than between unalike attributes of either option. Therefore, patterns of gaze transitions were consistent with choice probabilities and accuracies that indicate monkeys compared unintegrated attributes.

### OFC neurons encode attributes relative to the focus of gaze

Having quantified gaze behavior, we next asked whether OFC neurons encode the value of attended attributes or options. To do this, we aligned neural activity to the onset of pre-choice fixations, and used multiple linear regressions to determine how firing rates varied with the value of attributes relative to the focus of the subjects’ gaze. Each attribute of the choice was labeled as either the fixated attribute (e.g., sweetness A), the fixated attribute’s pair in the same option (e.g., probability A), the attribute in the unfixated option that is like the fixated one (e.g., sweetness B), or the other attribute in the unfixated option (e.g., probability B) (**Fig 4a**). If OFC neurons encode attended attributes only, then we expected neuron activity to be explained by the value of the fixated attribute alone, but if they encoded the integrated value of the attended option, then firing rates should be modulated by both the fixated attribute and its pair (green attributes in **Fig 4a**). On the other hand, if OFC neurons compare like attributes, we expected firing rates to encode the fixated attribute and the like attribute of the unfixated option (dark green and dark orange in **Fig 4a**). Since evidence suggests that gaze shifts early in a decision are primarily used for evaluation and later ones are confirmatory once a latent choice is identified (*46–48*), we included only the first and second fixations on trials with 2 or more pre-choice fixations (**Fig 3b**). Further, to focus on neural activity that shifts with the targets of gaze, we first regressed out activity related to static chosen and unchosen attributes (Methods).

**Figure 4.**
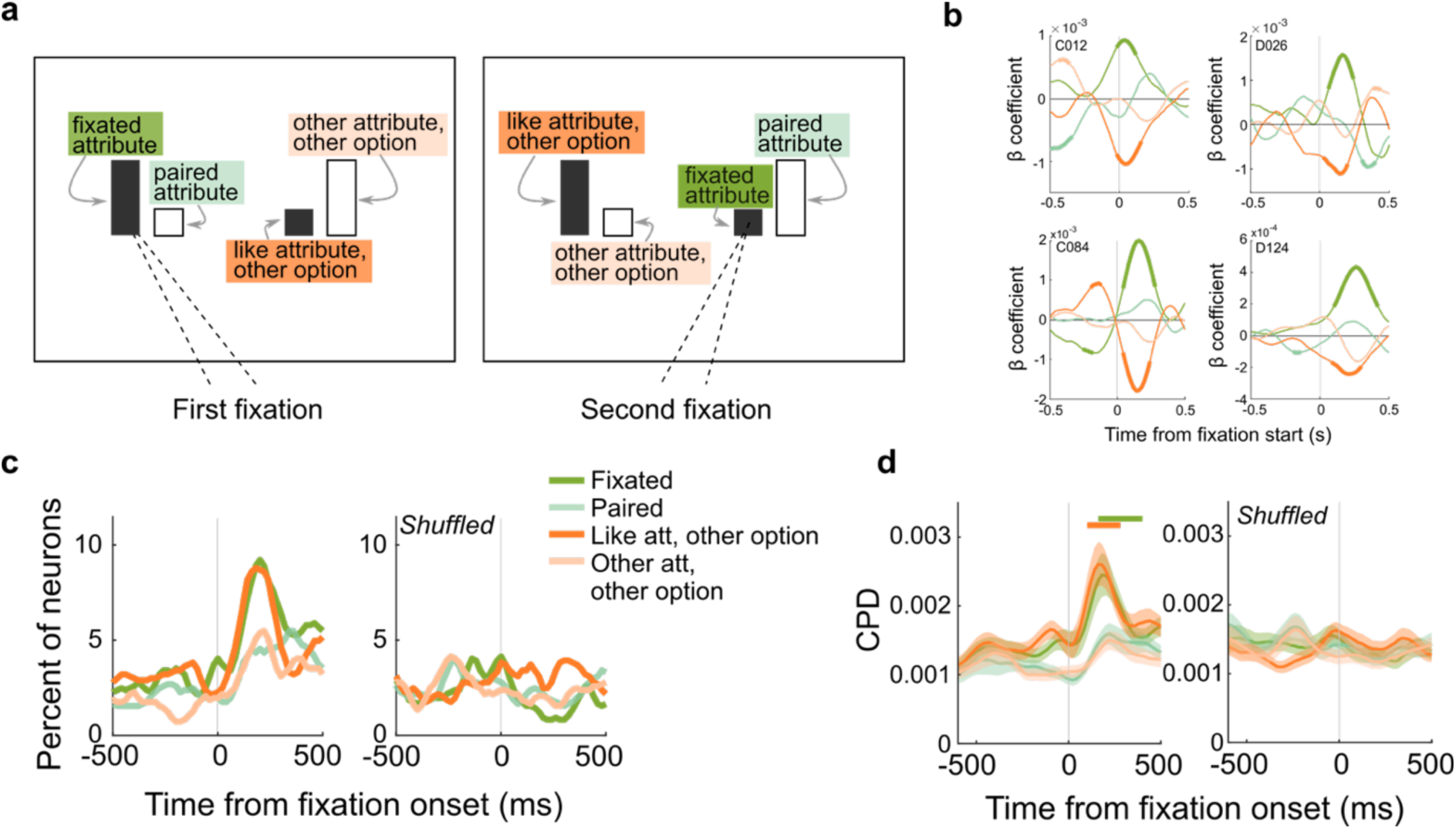
OFC neurons encode attributes relative to the focus of gaze. (**a**) Schematic showing attribute labels, depending on the monkeys’ targets of gaze for two example fixations. The first fixation is on the sweetness bar of the left option, and the second fixation is on the sweetness bar of the right option. (**b**) Four example neurons encoding attributes in an attention-based reference frame. Plots show beta coefficients from multiple regressions, with firing rates aligned to the onset of a fixation. Thick segments denote regions of significance (p≤0.05 x 3 consecutive time bins). Colors as in **a**. (**c**) Percent of neurons in intact (left) and shuffled data (right) that significantly encode each attribute. (**d**) CPDs for each attribute across all recorded neurons. Shading shows mean CPD +/− sem. CPDs were compared to shuffled controls with rank-sum tests. Bars: p ≤ 0.05 x 3 consecutive time bins.

Soon after directing gaze to an attribute, many OFC neurons encoded attribute values relative to the focus of the monkey’s gaze. In particular, neurons tended to encode the fixated attribute, often with a positive beta coefficient, and the like attribute of the unfixated option, often with a negative beta, as shown in example neurons in **Fig 4b**. This pattern, equivalent to a weighted subtraction, is consistent with competitive inhibition between an attended attribute and the matching attribute in the other option, and suggests that OFC neurons do carry out attribute-level comparisons. The comparison signal, however, appears to depend on directed attention, since the same signatures were not found in our previous analyses that did not take gaze into account. In the following sections, we quantify this tendency across our recorded population of OFC neurons.

Overall, there was a clear increase in neurons encoding values of both the fixated attribute and the like attribute of the other option soon after the start of a fixation (**Fig 4c**). At the same time, we did not find a change in the proportion of neurons encoding the other attributes on the screen, including the paired attribute in the same option that should be encoded if the neurons integrated option values. However, the proportion of neurons that reached our threshold for significance at any time was modest (≤10%), so we also calculated the unique variance accounted for by each of the attributes across all neurons in our data set, using the coefficient of partial determination (CPD). Similar to the thresholded results, there were peaks in variance explained by the fixated attribute and the like attribute of the other option soon after the start of a fixation (**Fig 4d**). Therefore, at the population level, there was increased encoding of like attribute values, relative to the focus of the monkeys’ gaze.

Before analyzing this effect further, we first ensured that our results were related specifically to directed gaze, and were not artifacts of the analyses. First, we ran a shuffling procedure in which the same attribute values were included in each trial, but their relationship to the monkey’s gaze was shuffled. In other words, we used the same residualized data, but attribute values were randomly designated the fixated attribute, paired attribute, like attribute of the other option, and the other attribute of the other option. As expected, shuffling removed the encoding that followed fixation onset in both thresholded data and CPDs (**Fig 4c-d**). Further, if only unfixated attributes were shuffled, then only encoding of the fixated attribute was found (**Fig S3g**).

Next, since monkeys tend to look at better attributes and chosen options first, we ensured that encoding patterns were not spurious effects of correlations between the monkeys’ gaze and the value of the choice attributes. To do this, we simulated neuron responses that encoded task variables but were agnostic to the fixation patterns the monkey generated, then tested whether the analyses we used produced spurious encoding of fixated attributes. We simulated neuron responses of six types, corresponding to the most common encoding patterns found in our data set: neurons encoding chosen sweetness only, chosen probability only, better sweetness only, better probability only, chosen value (i.e., both chosen sweetness and chosen probability), and non-selective neurons (**Fig 5a**). We simulated firing rates that reflected only the variable being tested, plus random noise, using actual trial variables (e.g., chosen sweetness/probability on each trial) from each session, and then tested the simulated neurons on actual patterns of fixations produced by the monkey on the same trials. Using this approach, we found <1.5% of simulated neurons of each type encoded attributes relative to the monkeys’ fixations (**Fig 5b-g**), consistent with the expected false alarm rate of the significance criterion we used (p≤0.01). Therefore, there was no evidence that fixation-dependent encoding arose from confounds in the behavior or analyses.

**Figure 5.**
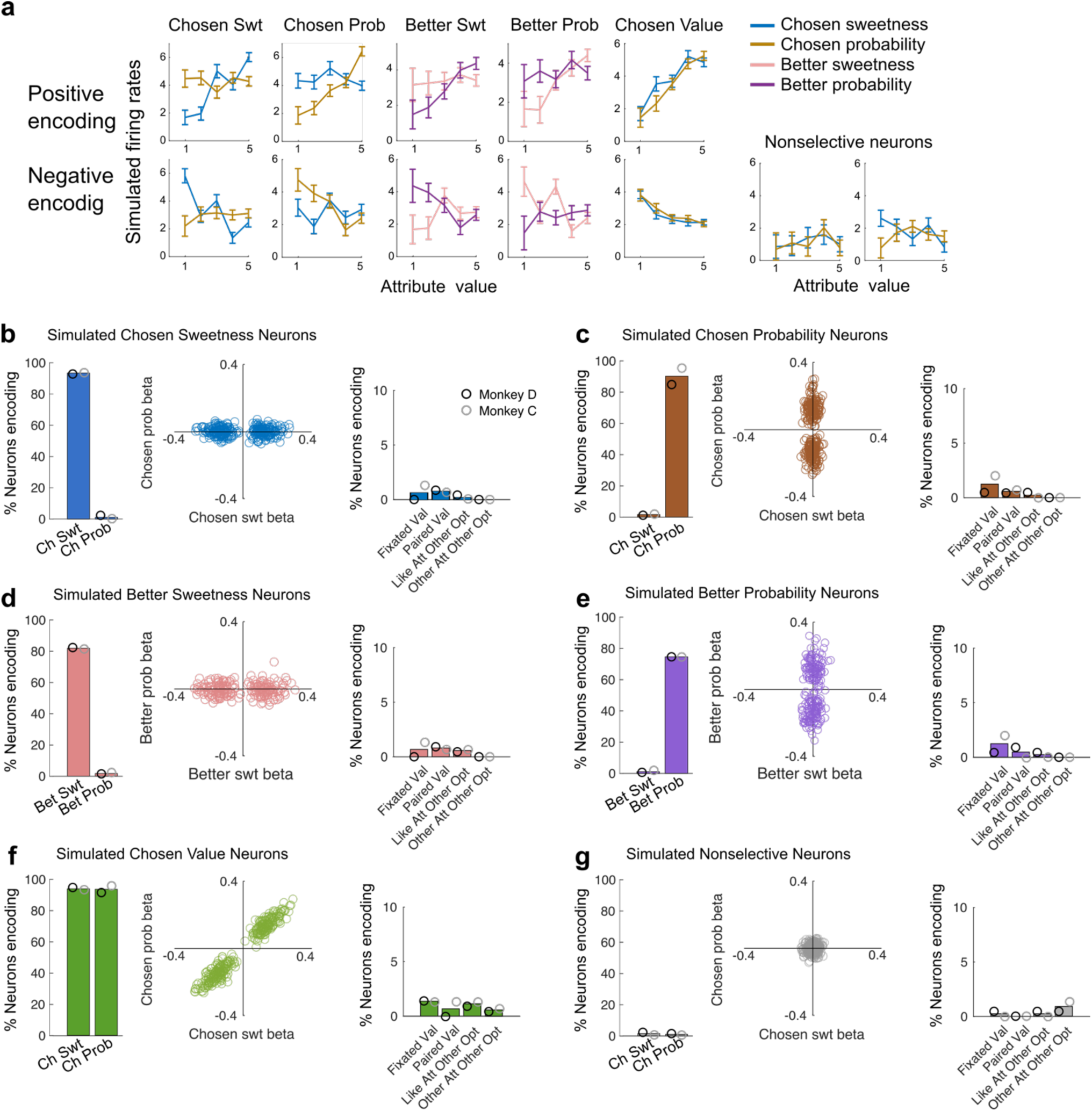
Simulated neurons encoding trial-wise attributes do not encode fixated attributes. (**a**) Two examples each of six types of simulated neurons. Average firing rates (+/− sem) of the simulated neurons are plotted at five values of each attribute. Plots show firing rates varying with chosen attributes or better attributes, depending on which type of encoding was simulated. For the first five types, the top row shows simulated neurons that positively encode the target attribute(s), and the bottom row shows negatively encoding simulated neurons. Nonselective neurons are neither positive nor negative. (**b**) The population of simulated neurons encoding chosen sweetness. The first plot shows the percent of neurons encoding chosen sweetness (Ch Swt) or chosen probability (Ch Prob) on mulitple linear regression as in the main data set. Significance = non-zero coefficients in the regression (p≤0.01). The second plot shows that beta coefficients for chosen sweetness and chosen probability produced the expected encoding pattern. The third plot shows the percent of neurons encoding fixation-related variables, after the simulated firing rates were residualized for trial-wise variables as in the main data set. Bars in the first and third panels are percents across both monkeys, black circles = Monkey D, gray circles = Monkey C. (**c-g**) The same as **b**, except for simulated encoding of chosen probability (**c**), better sweetness (**d**), better probability (**e**), chosen value (**f**), or no task-related variables (**g**).

Finally, encoding of fixated and like attributes occurred ∼200ms after the start of a fixation. Since each fixation lasted ∼200ms (**Fig S3a-b**), and monkeys tended to shift their gaze between like attributes, we assessed whether encoding assigned to the like attribute of the other option was actually a consequence of the subject moving their eyes to that attribute on the second fixation. To check this, we analyzed the first and second fixations separately. Following each fixation, we obtained similar results in both proportion of neurons encoding these variables and CPDs, each occurring approximately 200ms after the respective fixation (**Fig S3c-f**). Interestingly, on the second fixation only, there was also encoding of the two relevant attributes before the fixation was initiated. This was separate in time from post-fixation encoding, and can be attributed to the prior (first) fixation (**Fig S3e-f**). Together, these analyses confirm that OFC neurons did encode both the fixated attribute and the like attribute in the other option at the same time.

### OFC neurons jointly encode attributes

Next, we assessed whether different attributes tend to be jointly encoded by single OFC neurons, as in the example neurons in **Fig 4b**, or if the population effects resulted from different pools of neurons encoding a single attribute at a time. We considered four possible combinations of two attributes that might be encoded together: the fixated attribute and its pair, as expected if the neuron encoded the attended option’s value (**Fig 6a**, pink); the fixated attribute and the like attribute of the other option, as expected if like attributes are compared (**Fig 6a**, blue); the two like attributes that were not fixated (**Fig 6a**, brown); and the two attributes of the unfixated option (**Fig 6a**, purple). We found a small but significant proportion of neurons jointly encoding the fixated attribute and the like attribute of the other option, consistent with the possibility of within-attribute comparisons (**Fig 6b, S4b**). Interestingly, we also found a small number of neurons that jointly encoded the two like attributes that are not fixated, which may be related to the monkeys’ tendency to peripherally evaluate attributes. On the other hand, there was no tendency to jointly encode the two attributes of either the fixated or unfixated option, again arguing against integrated encoding of option values in this gaze-dependent reference frame. The same patterns were found when data were separated into first and second fixations or by monkey (**Fig S4c-d**).

**Figure 6.**
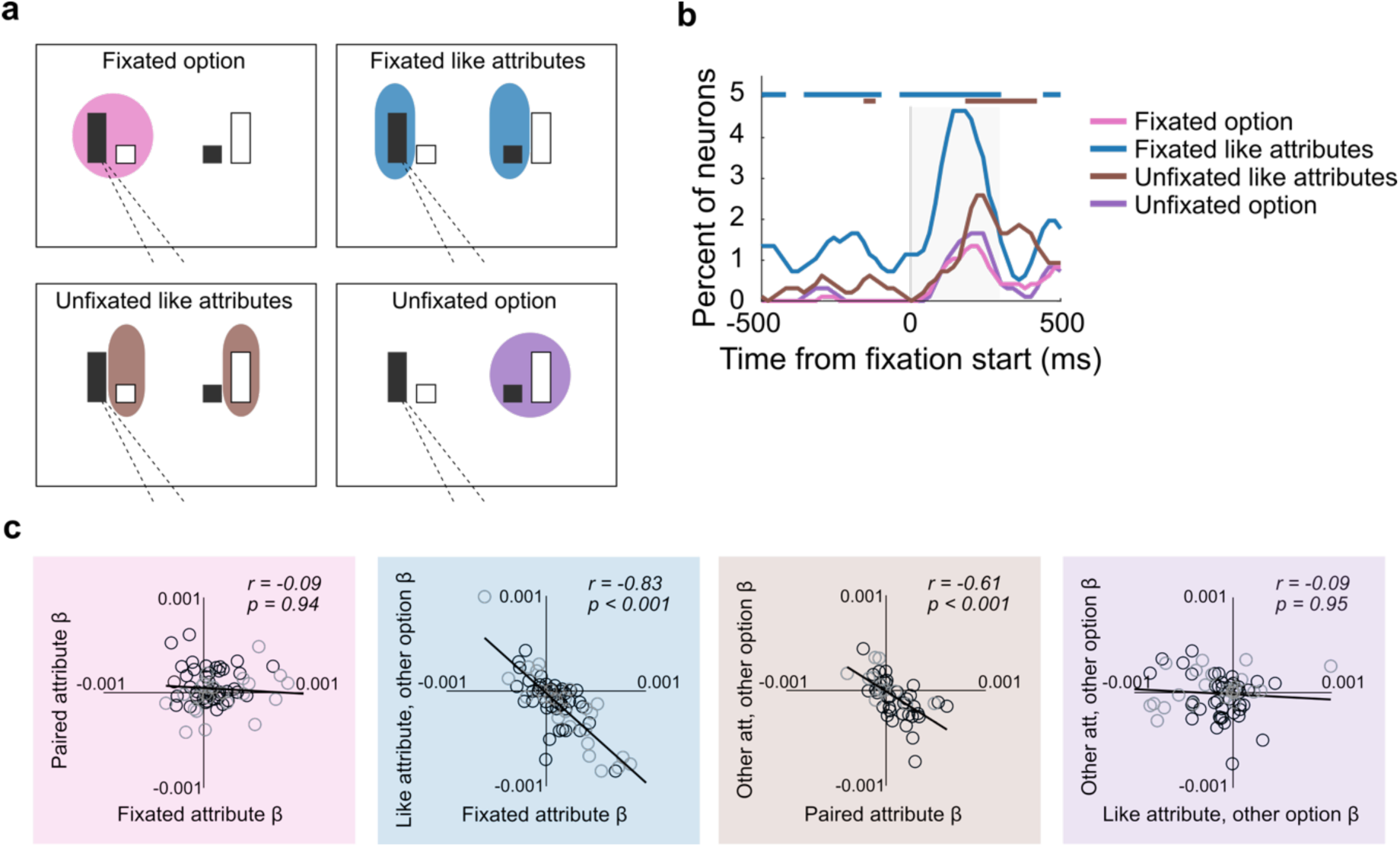
Neural signatures of attribute comparison in OFC. (**a**) Schematic of pairs of attributes that might be grouped in an attention-dependent reference frame. Each panel shows groupings for an example where the sweetness bar of the left option is fixated. (**b**) Percent of neurons jointly encoding each pair of attributes defined in **a.** Shuffled data are in Fig S4b. Bars = incidence of joint coding is greater than expected if the two attributes of a group were coded independently. To correct for each attribute belonging to two defined groups, significant joint encoding was defined by binomial test p ≤ 0.005 x 3 consecutive time bins. Shaded region = time windows analyzed in the scatter plots in **c**. (**c**) Scatter plots of mean beta coefficients of neurons with significance for one or both of the attributes in a group, color coded as in **a**. Statistics are from Pearson correlations, with p-values Bonferroni corrected for two comparisons each. Black circles = Monkey D, gray circles = Monkey C.

### Attribute comparison takes place in a gaze-based reference frame

Together, these results are consistent with the possibility that OFC neurons compute comparisons among like attributes during multi-attribute choice. In a final analysis, we tested for neurophysiological signatures of value comparison in this gaze-dependent reference frame. Our earlier analyses found no evidence for value comparisons, either among attributes or options (**Fig 2f-g**), but this could be because OFC computes comparisons in an attention-dependent reference frame and those analyses did not take gaze into account. Therefore, we tested whether values of the unfixated attribute or option antagonistically weight value encoding of the fixated attribute/option. To do this, we computed the average beta coefficient in a time window following an attribute fixation, and included neurons that reached significance for one or both regressors of interest at any time in that window.

Consistent with comparisons of like attribute values, we found an inverse relationship between the beta coefficients of the fixated attribute and the like attribute of the other option (**Fig 6c**, blue). In the same time window, there was no relationship between the two attributes of either the fixated or unfixated option (**Fig 6c**, pink or purple, respectively), indicating no evidence that OFC neurons encode integrated values in a gaze-dependent manner. Interestingly, there was a negative relationship among coefficients of the two like attributes that were not fixated (**Fig 6c**, brown), again likely arising from the subjects’ tendency to peripherally evaluate information. Since there was no relationship among attributes within the same option, the negative relationships among like attributes that were both fixated and not fixated could only arise if separate populations of neurons were modulated by each (fixated and not fixated). In sum, our results demonstrate a consistent neural signature of value comparisons that take place among like attributes, dependent on the monkeys’ current focus of gaze.

Overall, there was a bias among OFC neurons to encode the fixated attribute positively, meaning that firing rates increased with increasing value (**Fig 6c**). The like attribute of the other option, on the other hand, tended to be encoded negatively. Since the first gaze transition tended to shift between these two attributes (**Fig 3f**), the first fixated attribute had a high probability of subsequently becoming the like attribute of the unfixated option (**Fig 7a**). Therefore, we expected the beta coefficients to flip from encoding the first fixated attribute positively to negatively at the start of the second fixation, and vice versa for the like attribute of the other option. To test this, we ran a multiple regression on the firing rates aligned to the first and second fixations separately, but used regressors defined by the first fixation throughout. As predicted, ∼200ms after the first fixation, average coefficients across the recorded population were positive for the fixated attribute and negative for the like attribute of the unfixated option. Approximately 200ms later, however, the signs flipped (**Fig 7b**). Aligning the same data to the second fixation revealed that the inverted encoding pattern peaked approximately 200ms after the second fixation, and the same effect was apparent in each monkeys’ data individually (**Fig S4e**). Taken together, these patterns confirm predictions of a model in which OFC neurons inversely encode like attributes of multi-attribute options, in a reference frame defined by the monkey’s focus of gaze.

**Figure 7.**
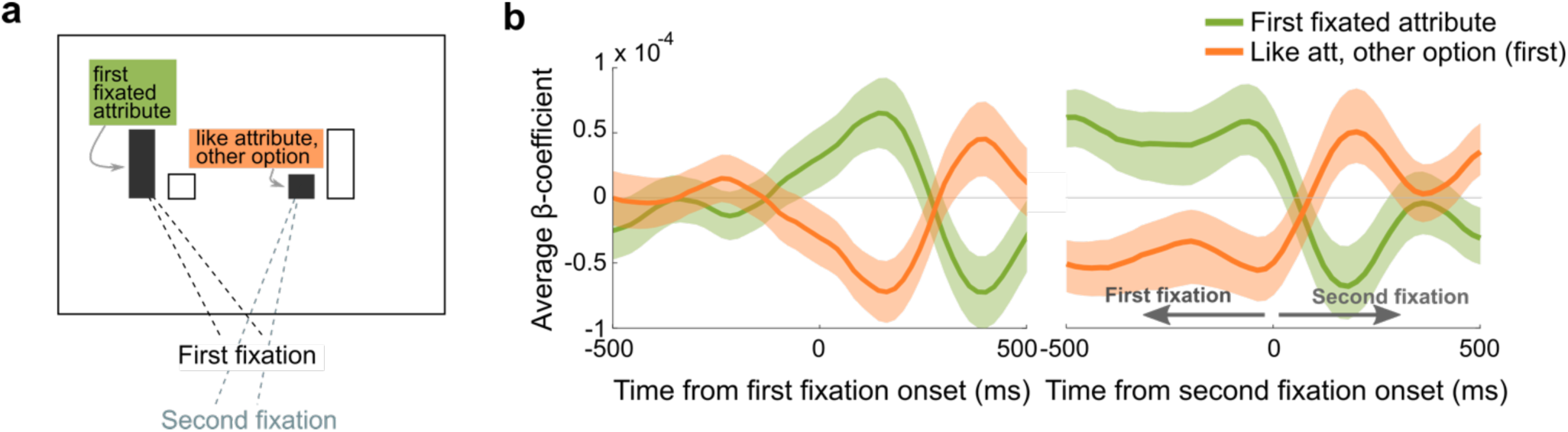
Attribute encoding is modulated by gaze shifts. (**a**) Schematic illustrating that a gaze transition between like attributes reverses the attribute designations. If the like attribute of the other option from the first fixation (orange) is fixated second, then it becomes the fixated attribute and the first fixated attribute (green) becomes the like attribute of the other option. (**b**) Average beta coefficients from trials with two or more pre-choice fixations. Firing rates are aligned to the first fixation (left) or the second fixation (right), with the same regression model in each. Shading shows the mean coefficient estimate across all neurons +/− sem.

## Discussion

Using a multi-attribute decision-making task, we found that OFC neurons encode and compare the values of like attributes, but not integrated option values. When neural responses were aligned to trial events, neurons predominantly encoded single attributes, initially relative to each other (better versus worse), and then relative to the choice (chosen versus unchosen), suggesting that OFC computes choices by comparing attribute values. However, value comparisons were not anchored to the monkeys’ choices or the attributes’ relative values, but were dynamically determined by the focus of the monkeys’ gaze. Since comparisons of integrated values have been reported outside of OFC (*4, 23–25*), our results support the proposal that multiple decision-related computations take place, potentially in parallel, throughout the brain (*49–52*). In OFC, this process is dynamic, dependent on the current target of attention, and focuses on the components that need to be weighed in order to make a decision.

As in previous studies, we defined value comparison signals as antagonistic effects of two cue values on the firing rate of an individual neuron, measured as inverse relationships between regression coefficients for the respective cues (*4, 23, 24, 26*). This is quantitatively related to a value difference, which has also been used to define potential value comparisons (*9, 14, 19, 20*), except the subtracted components have independent weights. This type of response indicates a competition between the cues, since firing rates are simultaneously driven up in proportion to one cue’s value and down in proportion to the other. The resulting neural responses correlate with the relative value of the cues, yet the underlying circuit mechanisms that produce this signature are still unclear. One suggested mechanism is mutual inhibition among value coding neurons (*4, 23, 27, 53*). From this view, activity in a pool of neurons selectively encoding the value of one cue (in our case an attended attribute), inhibits other neurons encoding a different cue (the like attribute of the unattended option), and vice versa. Since neuron activity depends on cue value, so does their inhibitory influence, resulting in an antagonistic firing rate code. While this is possible, the gaze-dependent nature of the comparisons we found presents a challenge for the mutual inhibition model. While some OFC neurons have reliable selectivity for attended cues (*22*), these would have to selectively inhibit neurons encoding the unattended sweetness when the monkey fixates a sweetness bar and the unattended probability when he fixates a probability bar, and dynamically shift inhibition targets as the subject gazes around the screen. Alternatively, a more general competitive inhibition process might produce the same effects (*21*), as could competition arising from limited resources such as attention. For example, if the monkeys covertly divide attention between compared attributes but allocate attention in proportion to value, then higher value alternatives would subtract more attention and ultimately produce an antagonistic effect on the attended item, as described here. Relative value coding of a similar pattern can also be produced by divisive normalization, a canonical computation that normalizes one neuron’s response by the activity of nearby neurons (*54*). Divisive normalization of value representations have been described in the lateral intraparietal cortex for cues in versus out of a receptive field (*55*), as well as in medial OFC for risky and safe options (*56*). Additional work is needed to test which candidate mechanism computes attention-dependent comparisons in OFC.

Many studies have attempted to isolate comparison signals by presenting options to subjects sequentially (*9, 18, 19, 21, 23, 24, 26*). In some regions such as vmPFC and ventral striatum, this has revealed comparisons of integrated value (*23, 24*), but in OFC, results have been less clear (*18, 21*). For instance, when options have a single attribute, OFC neurons primarily encode the first or second cue value rather than joint or antagonistic encoding that would indicate option comparison (*19*). However, another study found value difference signals in OFC by sequentially presenting options that varied in two attributes (*9*). Our results suggest that those comparisons may actually have been among component attributes, which correlate with the overall values the authors calculated, and that sequential presentation helped to control how the monkeys’ attention was allocated. Consistent with this, multi-attribute choices that do not use sequential presentation or take attention into consideration have not found strong evidence for value comparison signals (*14, 15, 20*).

The proposal that OFC neurons compare values in an attention-dependent reference frame is corroborated by a recent study that revealed attributes of a choice to the subject one at a time (*22*). Antagonistic value coding was found between current and previously attended information, though this was not clearly driven by comparisons of like attributes. In the case where two like attributes were shown first, the data aligned closely with our results, but when two attributes within the same option were shown first, their values were positively correlated, consistent with an integrated value signal (*22*). In contrast, our task presented all attributes simultaneously, and allowed monkeys to sample and compare information in any order. Although this results in more complicated behavior, it reveals more naturalistic information sampling and decision-making strategies. In this case, we found comparison signals in OFC exclusively between like attributes, and no tendency to integrate attributes within the attended or unattended option. In fact, we only evidence for potential integration among chosen (but not unchosen) attributes later in the choice suggesting that this signal reflects the result of a decision rather than the input to it. From this view, integration prior to choice, as found in previous tasks with sequential presentations (*9, 22*) may only occur when the other option’s information is not yet available, and might reflect the current best estimate of the expected outcome.

Our results are also consistent with the mounting evidence that value encoding in OFC is strongly dependent on attention. This includes both overt shifts of gaze (*39*) and covert shifts of attention (*40*). Moreover, attentional modulation of value encoding appears to be a more prominent feature of OFC activity compared to regions of the cingulate or lateral prefrontal cortex, where the location of attended items seems to be more relevant (*22*). At the same time, we found clear evidence that monkeys also used peripheral vision to evaluate options. Their first fixations were not directed randomly, but were more often to higher value attributes, implying that they rapidly and covertly evaluated components of the display to direct their eyes (*45*). It’s unclear whether the attribute comparisons we report depend on peripheral perception of attributes in the unfixated option, or if this comparison is based on a stored representation of other attributes’ values. The study that presented attributes one at a time removed them from view after sampling and still found comparison signals, suggesting that OFC can make comparisons to remembered value representations (*22*). However, another study found that removing unfixated options from view roughly doubled the size of attentional biases in choice behavior, indicating that peripheral vision does play a large role in choice computation (*57*). Whether attribute comparisons in OFC are enhanced or altered by the presence of other options in peripheral vision remains to be investigated.

Taken together, our results demonstrate that OFC is involved in evaluating and comparing component attributes of multi-attribute options. This leaves open the question of how decisions are made when options do not share common attributes. On one hand, these choices might involve different decision strategies. For instance, attributes could still be evaluated separately in an accept/reject fashion (*58–60*). On the other hand, these decisions might uniquely require the use of integrated values. To date, there is stronger evidence that vmPFC, rather than OFC, is involved in comparing integrated values (*23, 53, 56, 61–63*), and this process may also involve attention (*61*). Going forward, determining how attention is allocated in different choice scenarios and its impacts on neural coding of decision variables across regions will be important in understanding how the brain uses information to make optimal decisions.

## Acknowledgements

The authors thank Joey Charbonneau and Feng-Kuei Chiang for comments on the manuscript, and Fred Stoll for comments on the analyses. Funding support: R01MH134845 and a fellowship from the Pew Scholars Program in Biomedical Sciences to ELR, F31MH127901 to AQP.

## Author contributions

Conceptualization, ELR and AQP; Methodology, ELR and AQP; Software, ELR and AQP; Formal Analysis, ELR and AQP; Investigation, AQP; Data Curation, AQP; Writing – Original Draft, ELR; Writing – Review & Editing, ELR and AQP; Funding Acquisition, ELR and AQP.

The authors declare no conflicts of interest.

## Methods

### Subjects

Two adult male rhesus macaques (D and C) participated in the experiment. At the start of recording, they were 9 and 7 years old and weighed 10.0kg and 8.2kg respectively. Each was surgically implanted a titanium headpost and cranial recording chamber made of biocompatible plastic (Monkey D - Ultem, Jerry-Rig USA. Monkey C – PEEK, Crist Instruments). The chambers were centered on stereotaxic coordinates calculated from 3T MR images obtained of each subject’s brain. In Monkey C, the chamber was placed over the left hemisphere; in Monkey D, the right. All procedures were in accord with the National Institutes of Health guidelines and were approved by the Icahn School of Medicine at Mount Sinai Animal Care and Use Committee.

### Task and Behavior

Monkeys sat in a primate chair in a darkened testing chamber, head-fixed facing an 18-inch computer monitor positioned 17 inches away. MonkeyLogic software (*64, 65*) controlled the behavior interface. Eye position was recorded by an infrared eye tracker (ViewPoint Eyetracker USB-400, Arrington Systems) at a sampling rate of 400Hz. This signal was oversampled by MonkeyLogic at 500Hz.

Subjects were trained to perform a multi-attribute decision-making task. Behavior in this task has been previously reported (*37*), and the present data were collected from the same subjects upwards of two years later. Each trial began with the appearance of a fixation cue in the center of the screen. When the monkey gazed at the fixation point and simultaneously held a touch-sensitive bar for 550 ms, two (80% of trials) or three (20% of trials) choice options were displayed on the screen. Options were shown at 2 or 3 of 6 potential positions in a hexagonal arrangement around central fixation. Option positions were selected randomly, with the constraint that they were never in adjacent positions on the hexagon. Only 2-option trials were analyzed in the present study.

Each option was represented by a pair of two bars. The width of each bar was 2 degrees of visual angle, and the height varied from 2 to 10 degrees. The size of the left bar of each pair indicated the sweetness level (25, 50, 75, 100, or 125 mM sucrose solution) and the right bar the probability level (30, 40, 50, 60, or 70%). The colors of the bars indicated whether the size of the bar increased with increasing sweetness/probability level (direct mapping), or decreased with increasing sweetness/probability level (indirect mapping). Bar size and mapping varied randomly and independently on each trial.

While the monkey continued to hold the touch bar, he had up to 5 sec to freely view the images while gaze position was tracked. They could make a selection at any time by holding gaze on any part of the desired option and releasing the touch bar. This resulted in the unselected option disappearing and triggered the reward delivery immediately after an option selection. If no option was selected within 5 sec, or if the touch bar was released when gaze was not directed toward an option, this triggered a 5 sec timeout, during which no reward was delivered and the screen displayed a red background. When the timeout was complete, the next trial could be initiated. Incomplete trials were excluded from analysis. Inter-trial intervals were 1s.

Rewards consisted of 0.33mL of fluid delivered over 500ms for Monkey D, and 0.297mL delivered over 450ms for Monkey C. Amounts were titrated during pretraining to account for each subject’s relative weighting of sweetness and probability, and balance as closely as possible their subjective weighting of the two attributes.

### Behavior models

Choices in each session were fit with two potential models that varied only in how attribute values were combined. In the additive model (*Eq.1*), sweetness and probability were independently parameterized and linearly combined.

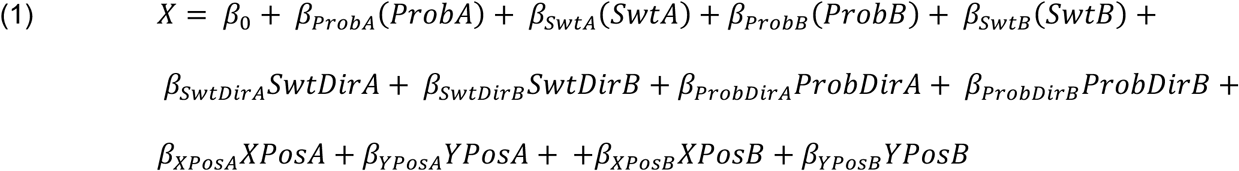

In the multiplicative model (*Eq. 2*), attributes were independently parameterized and multiplicatively combined.

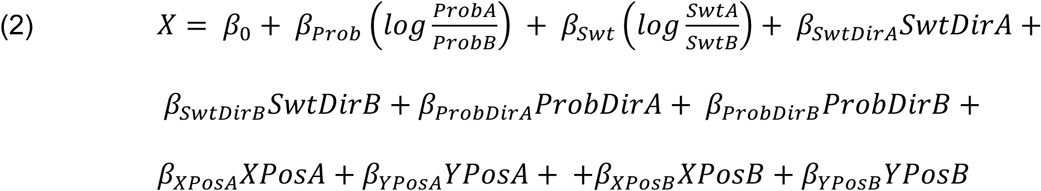

In both models, options were arbitrarily designated A and B. *ProbA* or *B* and *SwtA* or *B* are the ordinal magnitudes of probability and sweetness available in each option. *β_Prob/Swt_* are fitted weights of probability/sweetness attributes. *(Prob/Swt)Dir(A/B)* is the mapping for each attribute bar, coded as −1 or 1, with the corresponding coefficient (*β_(Prob/Swt)Dir(A/B)_*). *XPos(A/B)* is the x-position of option A or B on the task screen, and *YPos(A/B)* is its y-position. Since attributes within options were always proximal to each other and therefore correlated, only x and y coordinates of the center point of the overall option was used. *β_XPos(A/B)_* and *β_YPos(A/B)_* are the corresponding coefficients. *β*_0_ is a constant to capture any bias toward A or B. For both models, the probability of choosing an arbitrary option A was fit by generalized linear models with the logit link function. This is equivalent to a softmax decision rule (*66*) that probabilistically selects the option with the higher value (*Eq. 3*).

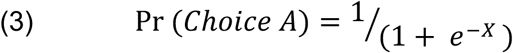

### Behavior model recovery

We simulated 100 sessions of 1000 choice trials each. As in choices presented to the monkeys, sweetness and probability magnitudes and mappings were randomly selected on each simulated trial, and each simulated option was assigned an X,Y position following the same constraints in the main task. Then, for each session, we used the parameters from the GLMs fit to the monkeys’ actual choices to predict a probability of selecting option A (Matlab function *predict.m*). This was done separately for additive and multiplicative models to obtain two vectors of trial-wise predictions, one for each model. For each set of predicted probabilities, we simulated choice outcomes on each trial by selecting option A with the probability *p*, and option B with probability 1-*p*, where *p* is the probability predicted by the model on that trial. Each simulated session was then fit by the same additive and multiplicative GLMs as the original behavior to determine whether the model comparison would reliably select the appropriate additive and multiplicative model when the generative model was known.

### Objective choice accuracies and condition-wise choice models

Among trials in which both attributes of one option were superior to both attributes of the other, accuracies were calculated as the proportion of trials per session in which the sweeter and more probable reward was selected, and one-way ANOVAs compared the average accuracies per session across the subsets of trials (as in **Fig 1d**). To model all choices in a condition (not just those with an objectively better option), trials with different mappings within options or within attributes were aggregated across all sessions.

From each condition, we created 100 bootstrapped samples of 400 trials and fit each set of trials with a simplified version of the additive model (*Eq. 4*).

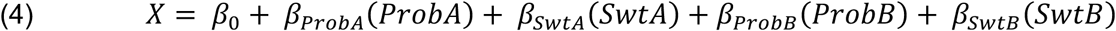

Beta coefficients for the same attribute of option A and B were approximately equal and oppositely signed, so we took the average absolute value as the estimated weight for each attribute. Weights were compared across mapping conditions with Wilcoxon rank-sum tests.

### Neural recording

Neurophysiological methods were similar to those reported previously (*67*). Data were collected with tungsten microelectrodes (FHC) and 16-contact linear arrays (Plexon) lowered through chronic craniotomies, and advanced manually using custom-built microdrives. Between 4 and 14 electrodes or 1 to 4 16-channel probes were lowered per session. Target regions were identified on previously obtained 3T MR images, as the cortex between medial and lateral orbital sulci. Our recording areas range from 35/42.7 mm to 28/32.7 mm AP for Monkeys D/C, relative to the interaural line. This included anterior regions putatively identified as Area 11 and posterior regions putatively identified as Area 13. After recording, electrode placements were reconstructed from MRIs, and any neurons recorded outside the target regions were removed from further analysis.

Neural signals were collected, digitized, and saved (Ripple Neural Systems Grapevine Processer). Putative spikes were captured at 30K and sorted into single and multi-units offline (Plexon OfflineSorter) and spike times were saved at 1kHz resolution. Only single units were analyzed in the present study, and any unit with an overall firing rate <1Hz was excluded from further analysis due to difficulty statistically characterizing their responses.

### Trial-aligned encoding models and model comparison

Prior to analysis, spike times were smoothed with a 150 ms boxcar. Smoothed firing was then aligned to the appearance of choice options and analyzed in 200 ms windows, stepped forward by 20 ms. Full time windows analyzed included 1s before until 1s after option onset, resulting in 90 overlapping time bins. In each time bin, *Eq. 5* was fit to the average firing rate. ProbCh and SwtCh are the ordinal magnitudes of the chosen probability and sweetness. ProbUCh and SwtUch are the ordinal magnitudes of the unchosen probability and sweetness. ProbDirCh and SwtDirCh are the mapping (direct or indirect) of the probability and sweetness attribute of the chosen option, and ProbDirUch and SwtDirUch are the mappings of the unchosen attributes. Each regression term was centered on zero before fitting the model.

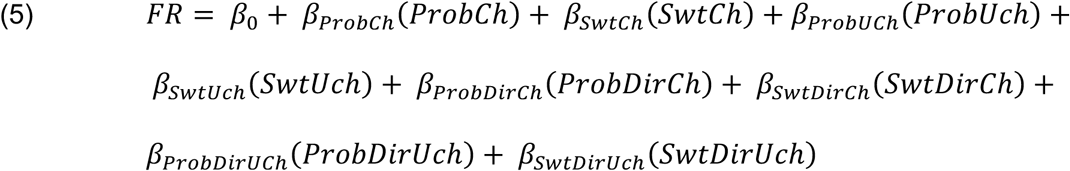

Significant encoding of a model term was defined as a non-zero beta coefficient (p≤0.01) for at least 3 consecutive time bins. We selected this criterion because it produced false discovery rates during the fixation epoch of approximately 5% or less for a single regression coefficient.

The same neuron firing rates in the same sliding windows were also fit with additional models. In each model, regressors were mean-centered before fitting. In the Better/Worse model (*Eq. 6*), ProbBet and SwtBet are the ordinal magnitudes of the higher value (better) probability and sweetness respectively and ProbWrs/SwtWrs are the ordinal magnitudes of the lower value (worse) probability and sweetness, as in **Fig 2b**. If an attribute had equal values in each option, then the same number was entered into each regressor. ProbDirBet and SwtDirBet are the mapping (direct or indirect) of the better probability and sweetness, and ProbDirWrs and SwtDirWrs are the mappings of the worse probability and sweetness.

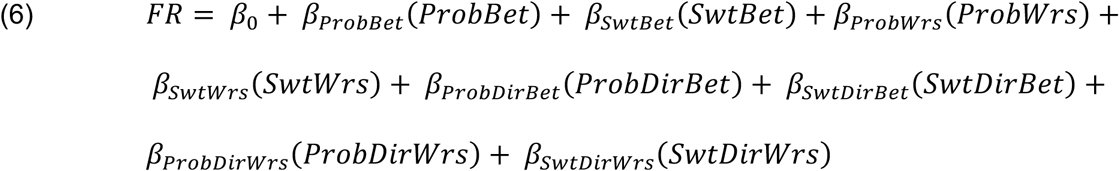

In the additive chosen value model (*Eq. 7*), attributes were first weighted by their relative importance in the additive behavior model for that session (*Eq. 1*). Since options A and B were arbitrarily designated in the behavior model, their weights were approximately equal and opposite. Therefore, we took the mean absolute value of the behavior coefficients for probability attributes (*β_BehProb_*) and multiplied this by trial-wise probabilities, the mean absolute value of behavior coefficients for sweetness attributes (*β_BehSwt_*) multiplied by trial-wise sweetnesses, then added these to obtain a weighted additive value for chosen and unchosen options. The model also included attribute mappings as in *Eq. 5*.

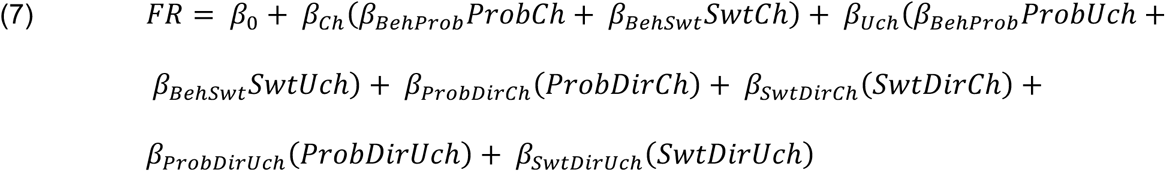

In the multiplicative chosen value model (*Eq. 8*), attributes were weighted by their relative importance in the multiplicative behavior model for that session (*Eq. 2*). This behavior model estimated one coefficient for the log ratio of probabilities (*β_Prob_*), and another for the log ratio of sweetnesses (*β_Swt_*). These were used as weights for each attribute, and attributes of chosen and unchosen options were multiplied to obtain integrated values. The model also included attribute mappings as in *Eq. 5*.

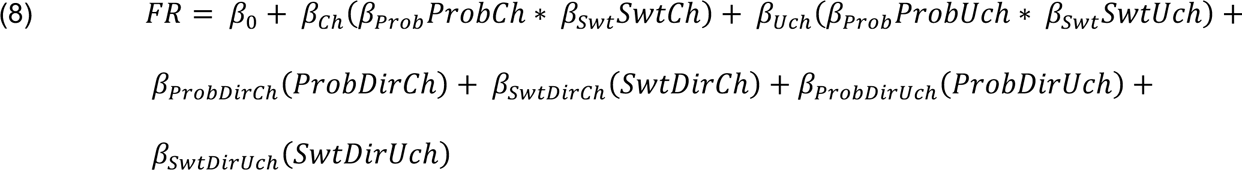

Finally, we considered the possibility that neurons encoded risk, or uncertainty, rather than probability (*68*). The risk model (*Eq. 9*) was identical to *Eq. 5*, except the ordinal levels of probability were replaced with ordinal levels of uncertainty, such that probabilities 1,2,3,4,5 were replaced with 1,2,3,2,1. RiskCh and RiskUch are chosen and unchosen risk attributes respectively.

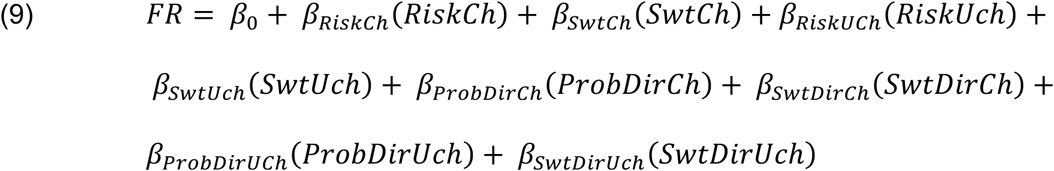

Each of the encoding models above were used to explain trial-wise variance in firing rates in 200 ms sliding windows. In each window, the total model significance was computed, as well as the Akaike information criterion (AIC). If the same model had a significance of p ≤ 0.01 for three consecutive time bins and had the lowest AIC of the models under consideration (*Eq. 5-7* in the main text, *Eq. 5-9* in Supplemental Data), then it was included as a neuron best fit by that model in the relevant time bins. Note that this means that the same neuron could be assigned to different models in different time bins. To determine whether single neurons initially coding better/worse attributes were more likely to code chosen/unchosen attributes later in the trial, we found the proportion of neurons assigned to the better/worse model in any time bin in the first 350ms after the choice appeared, and the proportion assigned to the chosen/unchosen model in any time bin between 350-700ms and tested whether the probability a neuron significantly encoding both of these was higher than expected if the two models were encoded independently (i.e., the joint probability of the two types of encoding). The rate of actual joint encoding was compared to this chance level with binomial tests.

### Gaze analyses

The EyeMMV toolbox (*69*) was used to define fixations from continuous eye movements. The minimum fixation duration was 50 ms to capture short fixations (*70*). The tolerance to include a gaze position in a fixation cluster (t1) and to include it in the calculation of the fixation cluster mean (t2) were t1=2 and t2=1. Only fixations that fell with defined regions around each attribute bar were included in the analyses. To capture fixations that fell on the bar edges, these regions included the full bar height plus 1.0° of visual angle around all sides except the side adjacent to the other attribute, which was 1.0° away. Here, an additional 0.5° buffer was used, so there was no unassigned space between the bars. The eye tracker was calibrated at the start of each session, and gaze data were further aligned in post-processing by centering eye traces on the mean x-y coordinates during the initial fixation window on each trial. Analyses excluded the fixation that coincided with release of the touch bar (i.e., the choice report). For each pre-choice fixation, EyeMMV was used to quantify fixation position (in x,y coordinates), start time, end time, and duration. Fixation start times were used to align neural data to fixations. Except where noted, the first and second fixations of trials with 2 or more pre-choice fixations were used.

Neural responses aligned to fixations and analyzed similarly to trial-aligned data. Firing rates were smoothed with 150 ms boxcar, averaged in sliding windows of 200 ms stepped forward by 20 ms, and analyzed with GLMs. To remove fixation-unaligned effects, we first fit each time window for each neuron with *Eq. 5*, then fit the residuals with *Eq. 10*, where FixVal and FixDir are the ordinal value and mapping direction of the fixated attribute (e.g., sweetness A), PairVal and PairDir are the ordinal value and mapping of the attribute that is the other component of the fixated option (e.g., probability A), LikeAttOtherOptVal and LikeAttOtherOptDir are the value and mapping of the like attribute of the unfixated option (e.g., sweetness B), and OtherAttOtherOptVal and OtherAttOtherOptDir are the value and mapping of the other attribute of the unfixated option (e.g., probability B). FixAtt indicates whether the currently fixated attribute is sweetness or probability, and XPosFix and YPosFix are the X and Y position coordinates of the fixation. Significant encoding was defined as p≤0.05 for three consecutive time bins. This criterion was selected because it resulted in <5% false discovery rates in shuffled data. Results were qualitatively similar if a stricter criterion was used (p≤0.01 for three consecutive time bins).

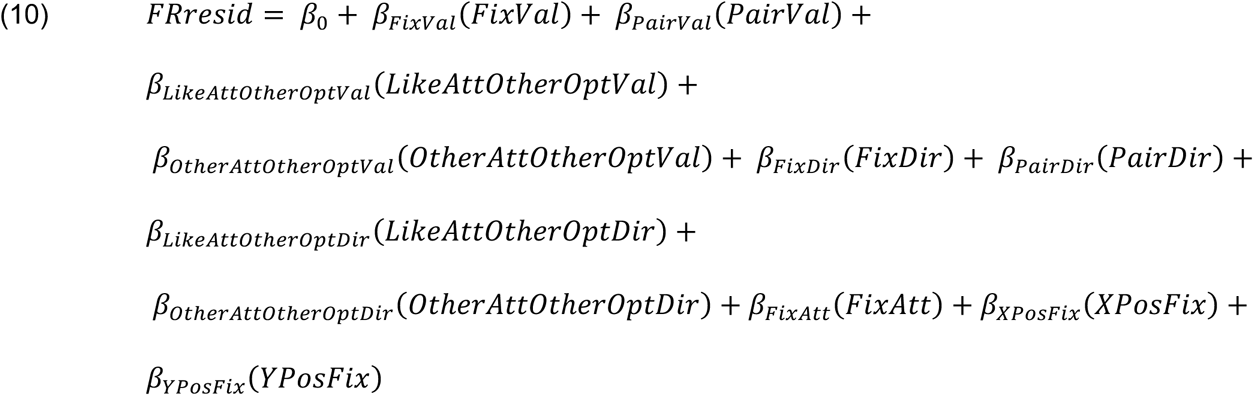

CPD quantifies the increase in unexplained variance obtained by removing one predictor from a multiple regression model, and therefore measures the unique variance explained by that predictor. CPDs were calculated for all neurons in the same sliding windows as above.

Joint encoding was defined as significant encoding (also p≤0.05 for three consecutive time bins) of two regressors in the same time bin. If the two regressors were encoded independently in a population, more coincident encoding would occur by chance as the prevalence of encoding either increased. This rate of chance joint encoding was calculated as the product of the proportions of neurons encoding either regressor, and the actual rate of coincident encoding was compared to chance with binomial tests. To quantify the tendency to code two regressors similarly or inversely, we averaged the regression coefficients for pairs of regressors from *Eq. 10* in time windows where the leading edge was >0ms and <300ms from the start of fixation. We included any neuron that reached significance for encoding of either of the regressors in one or more time windows in that range. Note that fixated and non-fixated attributes of both types had the same ordinal range in this analysis (1 to 5, mean centered to −2 to +2), so unequal range effects cannot explain beta anticorrelations in this task (*21*).

### Neuron simulations

Simulations were used to tested whether neurons that varied only with trial-wise variables could produce spurious encoding of fixated attributes, given the behavior patterns of our monkeys in this task. For simulated neurons, we used actual trial variables from each behavioral session, including sweetness and probability of the choice options, the monkey’s selection, and the first fixated attribute. We then simulated 5 neurons of each type (see **Fig 5**) per session, resulting in 215 and 150 simulated neurons of each type for Monkey D and C respectively. Each neuron was simulated as a single vector of trial-wise firing rates that was seeded with an integer response (1 to 5) that depended on the task variable being tested. Non-selective neurons were randomly assigned an integer 1 to 5. Chosen value responses were determined by binning the additive combination of chosen sweetness and probability into 5 bins. The integer indicated which of 5 partially overlapping, uniformly distributed ranges a new seed, representing an average firing rate, was drawn from. These ranges were chosen to match typical firing rates of OFC neurons, with the lowest value set to 0 and the highest 6.5. The average value of each range stepped up from 1 to 5, such that random draws from each range would produce average responses that were ordered 1<2<3<4<5. These responses were inverted (to 5<4<3<2<1) with a probability of 0.5 to simulate the tendency of OFC neurons to encode value both positively and negatively. We then added two sources of gaussian noise to each trial: one that was unique to each simulated neuron and one, on average 10% as large, that was shared across simulated neurons in the same session. Depending on the simulated variable, *Eq. 5* or *Eq. 6* was used to confirm that the simulated neurons had the desired encoding properties. We then tested whether the same analyses used to show fixation-related encoding (described above) would produce spurious results.

