## Supplementary material for "Attention-dependent attribute comparisons underlie multi-attribute decision-making in orbitofrontal cortex": Fig S

**Supplemental Figure S1.** Behavior is best fit by models with independent attributes.

**Supplemental Figure S2.** Heterogeneity of single unit encoding.

**Supplemental Figure S3.** OFC encoding of attributes in an attention-based reference frame.

**Supplemental Figure S4.** OFC neurons jointly and inversely encode attended attributes.

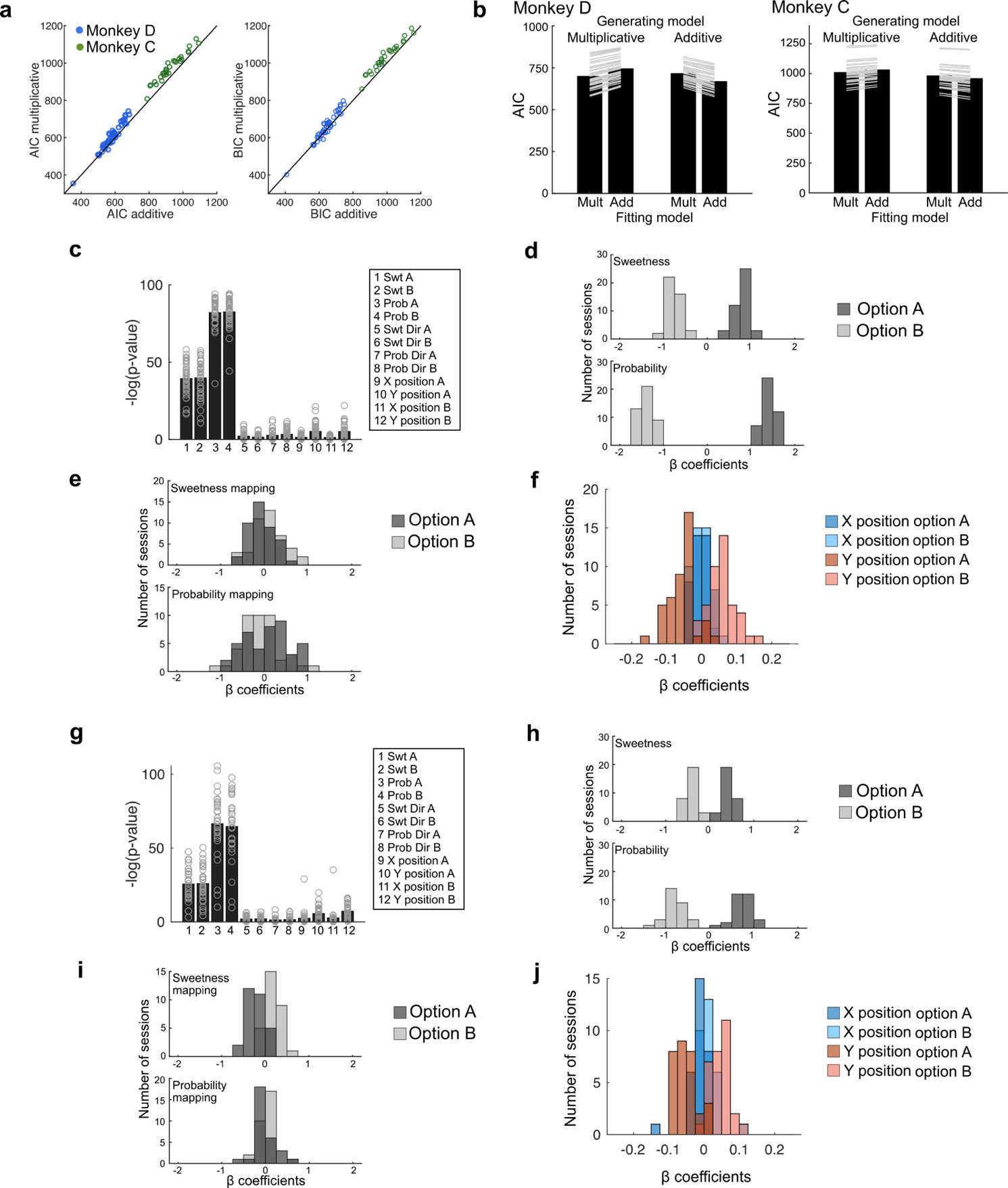

**Supplemental Figure S1.** Behavior is best fit by models with independent attributes. (**a**) AIC (left) and BIC (right) comparing behavior models in which attributes were combined additively or multiplicatively. Each point is a session, diagonal line = unity. In nearly every session, the additive model resulted in lower AIC and BIC values, indicating a better fit. (**b**) A model recovery procedure (see Methods) ensured that additive and multiplicative models produced choices that could be reliably differentiated. Data were simulated from either a multiplicative or additive model, then simulated data were fit with each model. AIC was used to compare fits to determine whether our analyses could reliably distinguish the correct generative model. For each animal (Monkey D left, Monkey C right), AICs were lower when the fitting model matched the generating model in every session. Bars show across-session means, lines show individual sessions. (**c - f**) GLM coefficients from additive models fit to choice behavior for Monkey D. (**c**) Significance of each coefficient estimate, shown as -log of the p-value calculated from the t-statistic testing for a non-zero coefficient. Bars show across-session means, circles show values from individual sessions. Coefficients labeled as in *Eq. 1*, where A and B refer to option A and B respectively, assigned arbitrarily in the GLM that predicted the probability of choosing option A. Attribute values had the largest effects on choices, and attribute mappings (labeled Dir) had almost no effects. (**d**) Histograms of coefficient estimates across sessions for the magnitude of each attribute. Positive/negative coefficients on option A/B indicate that the probability of choosing option A increases/decreases with the magnitude of that attribute. Larger deviations from 0 are indicative of more influence on the choice. (**e**) Histograms of coefficient estimates across sessions for the mapping of each attribute, as in **c**. (**f**) Histograms of coefficient estimates across sessions for the X and Y location of each option. Since attributes were always paired spatially, a single center location was used for each option. (**g - j**) Same as **c - f**, for Monkey C.

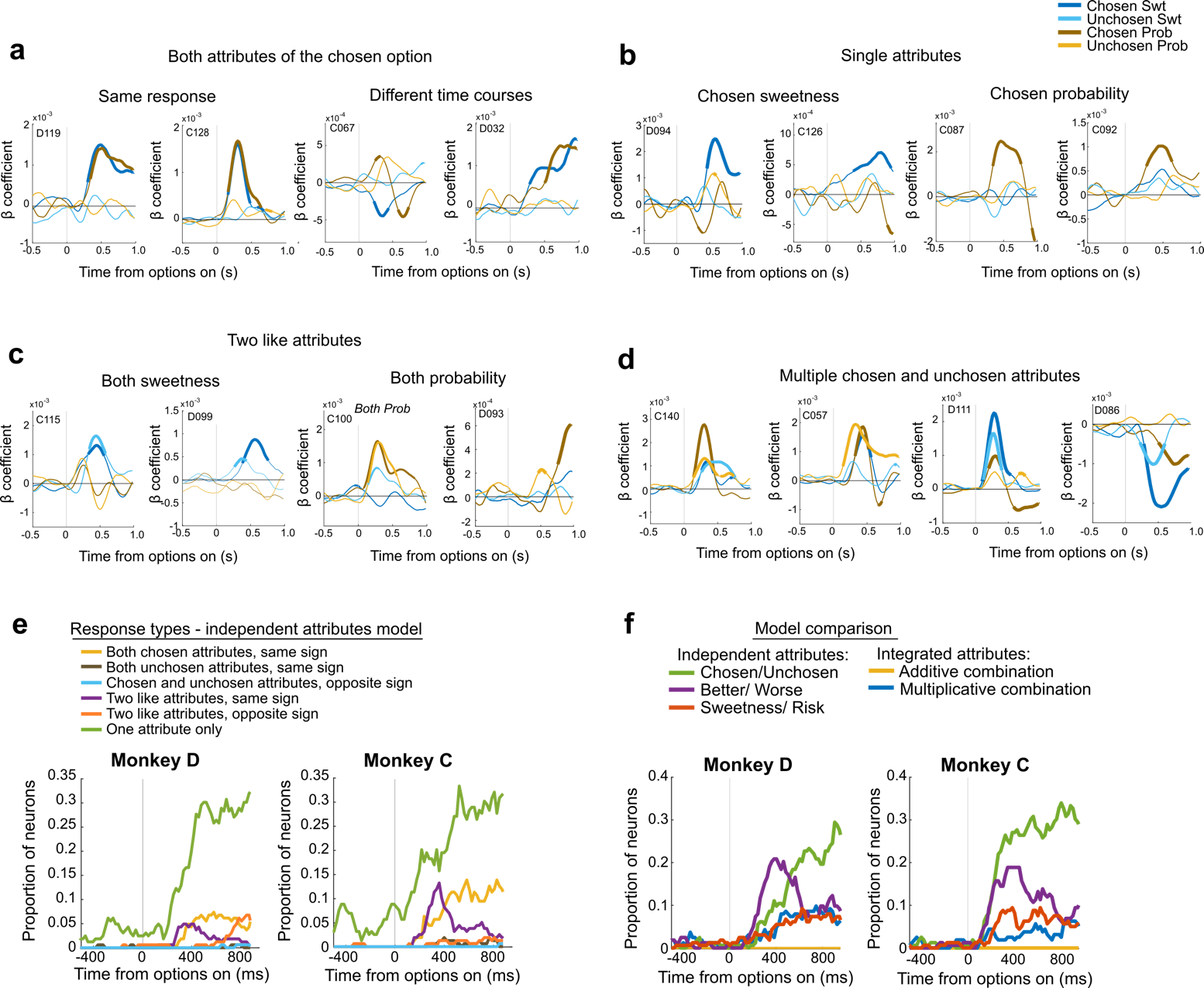

**Supplemental Figure S2.** Heterogeneity of single unit encoding. If single neurons in OFC construct option values, we expect to see neurons that encode both attributes of an option in a similar manner (i.e., with the same sign), and in approximately the proportion that each attribute is weighed in patterns of behavioral choices. (**a-d**) Examples of how many recorded neurons did not conform to these expectations. Thick lines = significant encoding (p≤0.01 x 3 consecutive time bins) in a sliding multiple regression model predicting neuron firing rates from chosen and unchosen attributes (*Eq. 5*). (**a**) Neurons encoding both attributes of the chosen option, but in manners inconsistent with this definition of an integrated value signal. The first two neurons encode the value of both attributes nearly identically, although behaviorally both subjects weighted probability slightly more than sweetness. These neurons may instead encode the ordinal rank of the chosen attributes. The second two neurons encode both attributes but with different time courses, also inconsistent with integration. (**b**) Further examples of neurons encoding only one attribute of the chosen option. (**c**) Further examples of neurons encoding both sweetnesses (chosen and unchosen) or both probabilities. (**d**) Examples of neurons with complex encoding patterns involving multiple chosen and unchosen attributes. (**e**) Tallies of neurons that exhibited different encoding patterns in the regression model that included chosen and unchosen attributes (*Eq. 5*), separately for each subject. The greatest proportion of neurons in both subjects encoded a single attribute at a time (green), and both subjects showed a small peak of neurons encoding two like attributes soon after the options appeared (purple). A small proportion of neurons encoding both attributes of the chosen option appeared slightly later (yellow). In contrast, there were almost no neurons encoding both attributes of the unchosen option, suggesting a lack of unchosen integrated value signals (blue). There were also very few neurons encoding two attributes with opposite signs (orange) or both attributes of the chosen option with opposite signs to the attributes of the unchosen option (teal), as expected if the comparisons between attributes or options were encoded. (**f**) Proportion of neurons in each subject best fit by different encoding models across time. For a neuron to be assigned to a model, it had to both predict significant variance in the neuron’s activity (full model p≤0.01 x 3 consecutive time bins) and provide the best fit as assessed by AIC. Here, we tested two additional models to the figure in the main text. In the first (red), chosen and unchosen probability was replaced with chosen and unchosen risk (or uncertainty), which was highest at 50% probability and lowest at both the lowest and highest probability. The second (blue) was a variant of the integrated value model, in which option values were computed as the multiplicative combination of weighted sweetness and probability, where the weights were determined by the attribute weights in the multiplicative behavior model for that session. Overall, the most commonly selected models were better/worse attribute values early in the choice, and chosen/unchosen attribute values later.

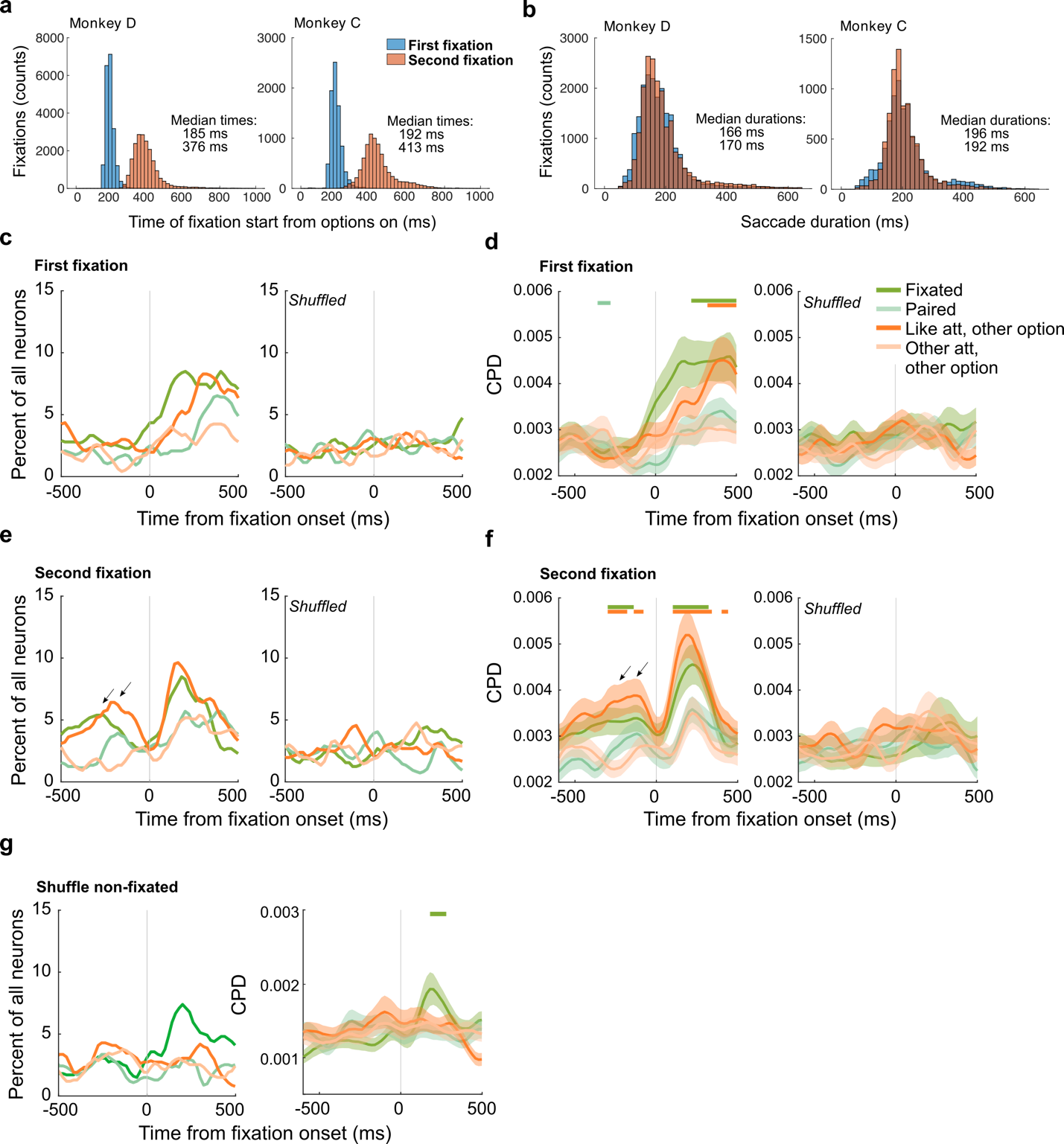

**Supplemental Figure S3.** OFC encoding of attributes in an attention-based reference frame. (**a**) Histograms of the time that the first (blue) and second (red) fixation begins, relative to the time that the choice options are presented, plotted separately by monkey. (**b**) Histograms of the duration of the first (blue) and second (red) fixations, plotted separately by monkey. (**c-d**) Percent of neurons that significantly encode each attribute (**c**) and CPD across all neurons (**d**) in intact (left panels) and shuffled data (right panels) from only the first fixation of trials with 2 or more pre-choice fixations (i.e., the trials analyzed in the main results). Line plots in **d** are the mean, shading is +/-sem. Bars indicate time windows where CPD is significantly greater than shuffled data (p≤0.05 x 3 consecutive time bins). (**e-f**) same as **c-d**, except for the only the second fixation of the same trials. Note that there are two epochs of significant encoding, one occurring after the second fixation begins, and one occurring before the start of the second fixation (arrows). The latter overlaps in time with the first fixation on these trials. (**g**) Plots as in **c-d**, of the same data as shown in the main results, except the values of the 3 unfixated attributes were randomly assigned to Paired, Like attribute other option, or Other attribute other option. Since only the fixated attribute assignment was intact, this was the only encoding recovered above chance.

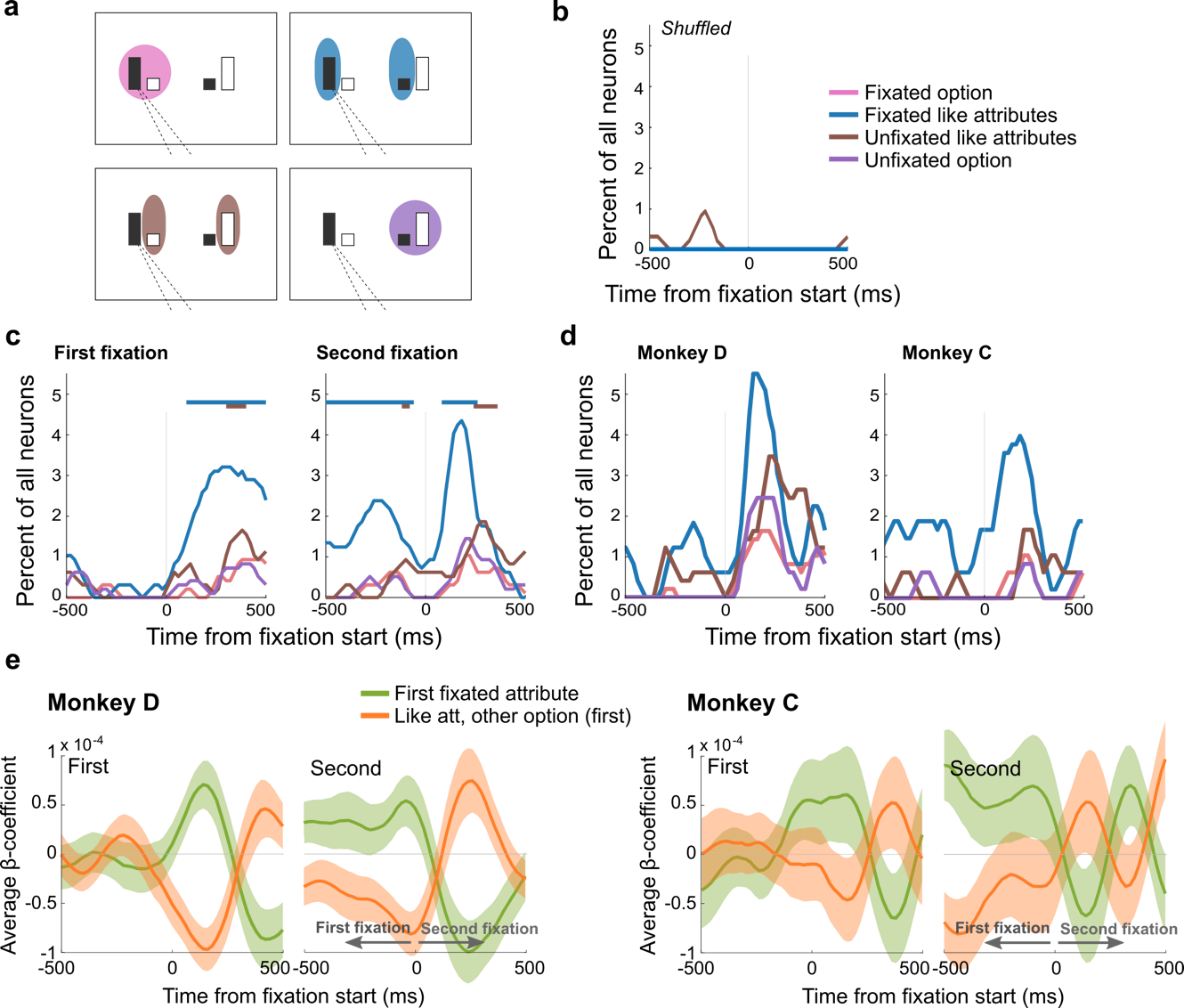

**Supplemental Figure S4.** OFC neurons jointly and inversely encode attended attributes. (**a**) The same schematic as in the main paper, showing pairs of attributes that were assessed for joint encoding, reproduced here for reference. (**b**) Joint encoding in the same data as in the main results, except with fixation assignments shuffled. (**c**) Joint encoding of attributes computed separately on the first (left) or second (right) fixation of trials with 2 or more pre-choice fixations. Two-sided binomial tests were performed on each sliding window to compare the incidence of joint coding to chance, which was defined as the joint probability of encoding either attribute of a pair. Bars indicate time windows that reached significance (p≤0.005 x 3 consecutive time bins). Note that there was significant joint encoding both following and prior to the second fixation, the latter overlaps with the time of the first fixation. (**d**) Joint encoding of attributes computed separately by monkey. The most common joint encoding in both animals were the fixated attribute and the like attribute of the other option (blue). (**e**) Average beta coefficients, separated by monkey, from multiple regressions of fixation-related attribute values. Regressions were performed separately for the first and second fixation on trials with two or more pre-choice fixations. The same regressors, with variables defined by the first fixation, were used across both fixations. As in the main results, both animals separately showed that the average coefficients tended to be positive for the fixated attribute and negative for the like attribute of the other option (green and orange respectively, at times after 0 on the first fixation and before 0 on the second fixation). Because monkeys tended to shift their gaze among like attributes, the second fixation was frequently directed to the attribute that was previously defined as the like attribute of the other option. Consistent with this, the average coefficients for that attribute became positive after the second fixation (orange), while the previously fixated attribute (green) became negative. Shading = +/- sem.
